# ACE2 Homo-dimerization, Human Genomic variants and Interaction of Host Proteins Explain High Population Specific Differences in Outcomes of COVID19

**DOI:** 10.1101/2020.04.24.050534

**Authors:** Swarkar Sharma, Inderpal Singh, Shazia Haider, Md. Zubbair Malik, Kalaiarasan Ponnusamy, Ekta Rai

**Affiliations:** Human Genetics Research Group, School of Biotechnology, Shri Mata Vaishno Devi University, Katra, Jammu and Kashmir, India; Bioinfores Pvt. Ltd., R. S. Pura, Jammu, Jammu and Kashmir, India; Department of Biotechnology, Jaypee Institute of Information Technology, Noida, Sector-62, Uttar Pradesh, India; School of Computational and Integrative Sciences, Jawaharlal Nehru University, New Delhi, India; School of Biotechnology, Jawaharlal Nehru University, New Delhi, India

## Abstract

Severe acute respiratory syndrome coronavirus 2 (SARS-CoV-2) is a positive single-stranded RNA virus that causes a highly contagious Corona Virus Disease (COVID19). Entry of SARS-CoV-2 in human cells depends on binding of the viral spike (S) proteins to cellular receptor Angiotensin-converting enzyme 2 (ACE2) and on S-protein priming by host cell serine protease TMPRSS2. Recently, COVID19 has been declared pandemic by World Health Organization (WHO) yet high differences in disease outcomes across countries have been seen. We provide evidences to explain these population-level differences. One of the key factors of entry of the virus in host cells presumably is because of differential interaction of viral proteins with host cell proteins due to different genetic backgrounds. Based on our findings, we conclude that a higher expression of *ACE2* is facilitated by natural variations, acting as Expression quantitative trait loci (eQTLs), with different frequencies in different populations. We suggest that high expression of ACE2 results in homo-dimerization, proving disadvantageous for TMPRSS2 mediated cleavage of ACE2; whereas, the monomeric ACE2 has higher preferential binding with SARS-CoV-2 S-Protein vis-a-vis its dimerized counterpart. Further, eQTLs in *TMPRSS2* and natural structural variations in the gene may also result in differential outcomes towards priming of viral S-protein, a critical step for entry of the Virus in host cells. In addition, we suggest that several key host genes, like *SLC6A19, ADAM17, RPS6, HNRNPA1, SUMO1, NACA, BTF3* and some other proteases as Cathepsins, might have a critical role. To conclude, understanding population specific differences in these genes may help in developing appropriate management strategies for COVID19 with better therapeutic interventions.

## Introduction

The recent emergence of corona virus disease (COVID19) caused by SARS-CoV-2 has resulted in >4.1 Million infections and >285 thousands deaths worldwide so far, and the numbers are increasing exponentially [https://covid19.who.int/]. SARS-CoV-2 is reported to be originated in bats (Zhou et al., 2020) and transmitted to humans via unknown intermediate host. However, its origin is still being questioned time and again. With the declaration of SARS-CoV-2 as pandemic by WHO, extensive research worldwide has been carried out. It has been established that Human ACE2 mediates SARS-CoV-2 entry into cells through its Spike (S) Protein, which primarily makes entry to host body through respiratory tract with nasal epithelial cells as potential initial infection site (Sungnak et al., 2020). ACE2 is a functional receptor on human cells for this newly originated coronavirus (Walls et al., 2020) with a higher affinity than the severe acute respiratory syndrome coronavirus (SARS-CoV) originated in 2002 (Wan et al., 2020). However, no substantial evidence exists about the higher expression of ACE2 being primarily associated with the degree of infection (Kuster et al., 2020). Additionally, COVID19 lethality is mostly driven by the extent of underlying lung injury; whereas, a negative correlation has been reported between ACE2 expression and lung injury (Imai et al., 2005). A recent report also suggests an inhibition of SARS-CoV-2 by human recombinant soluble (hrs) ACE2 (Monteil et al., 2020), making it an interesting question to explore.

High differences in clinical outcomes across countries have been [https://covid19.who.int/] noted which demonstrate that neither all people who are exposed to SARS-CoV-2 develop infection nor all infected patients end up in severe respiratory illness (Guan et al., 2020), which cannot be explained by immunity alone (Shi et al., 2020). This leads one to hypothesize about differential genetic susceptibility to COVID19 and virulence of SARS-CoV-2 in different populations (Kaiser, 2020). Efforts are being made to gain a better understanding of this disease and even propose prophylactic-hypothetical role of Bacillus Calmette–Guérin (BCG) vaccine, a vaccine primarily used against tuberculosis (TB), for reduced morbidity and mortality for COVID19 in human population (Miller et al., 2020). Due to extensive contemporary research, evidences in favour (Stawiski et al., 2020) as well as against (Cao et al., 2020) the existence of SARS-CoV-2 S-protein binding-resistant ACE2 natural variants, in different populations, have been found recently. In addition, studies highlighting role of eQTLs in *ACE2* expression, resulting in potential differential COVID19 fatality (Cao et al., 2020; Chen et al., 2020a), are pouring in. Where most such studies are targeting only natural variations in ACE2 gene as SARS-CoV-2 differential susceptibility factor, recent evidences suggest that additional host proteins like cellular serine protease TMPRSS2 act as co-factors and are critical for efficient cellular infection by SARS-CoV-2 (Hoffmann et al., 2020). Further, it is equally important to consider that rare functional variants, with uncertain consequences, may not explain large-scale population level differential clinical outcomes.

This highlights the importance of identification of other potential co-factors and underlying mechanisms these genes could be involved in. At the same time, understanding the interactions of these host proteins with SARS-CoV-2 along with ACE2 may explain many of the unanswered questions. Further evaluation of these host genes, and exploring their natural occurring variants, along with their expression patterns, may also shed some light on better conceptual framework of differential susceptibility to COVID19 and the virulence of SARS-CoV-2. In the present study, we have tried to exploit existing literature to understand mechanisms of correlations, of several relevant host genes, including ACE2, which interact specifically with some prominent viral proteins, using *in-silico* approaches to understand the role of these as factors responsible for population level differences.

## Methodology

As SARS-CoV-2 is primarily a respiratory pathogen, the present study was conducted to understand mechanisms of its entry to cells in lungs, and we believe it could be extrapolated to other respiratory tract tissues, to explore potential factors that are responsible for differential outcomes in respiratory illness. To begin with, we started with the most studied host protein ACE2 and tried to understand its interaction with SARS-CoV-2 S-Protein followed by adding other interacting proteins. We further added layers of other methods to have a better understanding of the potential causes and outcomes.

### Molecular Dynamics (MD) Simulations

Recently submitted experimental structure (6M17.pdb) of ACE2 dimer bound to two B0AT1 (coded by *SLC6A19)* and two receptor-binding domains (RBD) of S1 of SARS-CoV-2 Spike protein (S-protein), with overall resolution of 2.90 Angstrom and SARS-CoV2 Spike Glycoprotein with RBD domain in Up conformation (6VSB.pdb) were downloaded from the Protein Data Bank for structural visualization and dynamics analysis (Wrapp et al., 2020; Yan and Zhang, 2020). This structure was corrected for missing amino acids, side chains, missing hydrogen atoms, disulphide bonds etc. It was also optimized for pKa corresponding to physiological pH 7.2 and energy minimized to correct steric clashes using the Protein Preparation Wizard of Schrodinger Maestro (Madhavi Sastry et al., 2013). Predefined solvation model TIP3P was used and overall neutrality of the system was maintained by addition of Na+ and Cl-counter ions (Mark and Nilsson, 2001). Physiological salt concentration of 0.15 Molar was generated through addition of NaCl. Periodic boundary condition of 10 Angstrom was set using the System Builder Tool of Desmond software(2006). Total two MD simulations, each 10 nanosecond long, were conducted. In one of the simulations, six chains i.e. ACE2 dimer with each monomer bound to a B0AT1 and RBD domain was simulated, where the resulting solvated system consisted of 424,847 atoms. For the other simulation, monomeric ACE2 bound to viral RBD consisted of 174,900 atoms. Root mean square fluctuation (RMSF) and principal component analysis were done through R based BIO3D module (Grant et al., 2006). MMGBSA free energy of binding between proteins was calculated through Prime Software of Schrodinger Suite (Jacobson et al., 2002). Structural visualizations and images were traced using pyMOL and VMD (Jacobson et al., 2002) (Humphrey et al., 1996).

### Gene Expression in Alternate Transcripts and Regulation Analyses

Genotype-Tissue Expression (GTEx) portal at https://gtexportal.org/ was used to explore various gene expression profiles. By Tissue, Multigene Query as well as transcript browser was used to understand expression of splice variants and exons in Lungs. eQTLs were viewed by GTEx IGV Browser as well as GTEx Locus Browser was used to plot gene specific eQTLs. eQTLs and other gene regulatory information, splice variants, and ESTs were also explored through UCSC genome browser, https://genome.ucsc.edu/ with build GRCh37/hg19. Table browser was used to interact with various datasets to look for overlaps and filter outcomes. Retrieved information was saved in files as well as plotted on UCSC browser as custom tracks. Comparison of RNA expression and Protein expression, mainly in lungs, was done at the Human Protein Atlas (Uhlen et al., 2015) at www.proteinatlas.org. HaploReg v4.1 (https://pubs.broadinstitute.org/mammals/haploreg/haploreg.php) was used to annotate effect of noncoding genome variants of regulation motifs.

### Analyses of Allele frequency distribution of Variants in different population Groups

Genetic data for global population groups was explored through the GnomAD portal at https://gnomad.broadinstitute.org/, as well as dbSNP database of NCBI at https://www.ncbi.nlm.nih.gov/snp. Gene specific SNPs data was also retrieved from 1000 genomes (1000G) Phase 3 data set available through https://www.ncbi.nlm.nih.gov/variation/tools/1000genomes as per Homo sapiens:GRCh37.p13 (GCF_000001405.25). Data for the population groups belonging to five super population groups [African (AFR), Ad mixed American (AMR), East Asian (EAS), European (EUR) and South Asian (SAS)] was analysed. The following sub population groups were studied. AFR: African Caribbeans in Barbados (ACB), Americans of African Ancestry in SW USA (ASW), Esan in Nigeria (ESN), Gambian in Western Divisions in the Gambia (GWD), Luhya in Webeye, Kenya (LWK), Mende in Sierra Leone (MSL), Yoruba in Ibadan, Nigeria (YRI). AMR: Colombians from Medellin, Colombia (CLM), Mexican Ancestry from Los Angeles USA (MXL), Peruvians from Lima, Peru (PEL), Puerto Ricans from Puerto Rico (PUR). EAS: Chinese Dai in Xishuangbanna, China (CDX), Han Chinese in Beijing, China (CHB), Southern Han Chinese (CHS), Japanese in Tokyo, Japan (JPT), Kinh in Ho Chi Minh City, Vietnam (KHV). EUR: Utah Residents (CEPH) with Northern and Western European Ancestry (CEU), Finnish in Finland (FIN), British in England and Scotland (GBR), Iberian Population in Spain (IBS), Toscani in Italia (TSI). SAS: Bengali from Bangladesh (BEB), Gujarati Indian from Houston, Texas (GIH), Indian Telugu from the UK (ITU), Punjabi from Lahore, Pakistan (PJL) and Sri Lankan Tamil from the UK (STU).

LDproxy tools of LDlink 4.0.3 web-based suite (https://ldlink.nci.nih.gov/) was used to map proxy variants in strong LD and with putatively functional role. LDpop tool of the LDlink 4.0.3 was used for geographically annotating allele frequencies in 1000G populations; alternatively also done by a web-based tool Data wrapper (https://www.datawrapper.de/) in some of the figures.

### Computational structural analysis on SARS-CoV-2 S-protein and TMPRSS2

TMPRSS2 has been recently characterized as a critical component for cell entry by SARS-CoV-2 (Hoffmann et al., 2020). To understand its interactions, the protein sequence of surface glycoprotein (YP_009724390.1) of SARS CoV-2 and two transcripts of TMPRSS2 (NP_005647.3 and NP_001128571.1) protein of homo sapiens, were retrieved from the NCBI-Protein database. Pairwise sequence alignment of TMPRSS2 isoforms was carried out by Clustal Omega tool (Sievers et al., 2011). The protein domain information and transcript variation were retrieved from UniProt (UniProt, 2019), Prosite (Sigrist et al., 2013), Pfam (El-Gebali et al., 2019) and ENSEMBL (Chen et al., 2010), respectively. The homology model of the S-protein and TMPRSS2 were constructed using Swiss model (Waterhouse et al., 2018), whereas 3D structure of the ACE2 was retrieved from the PDB database (PDB ID: 6M17). These structures were energy minimized by the Chiron energy minimization server (Ramachandran et al., 2011). The binding site residues of the proteins retrieved from Uniport and literature. The mutant structure of the TMPRSS2 protein was generated using WHATIF server (Chinea et al., 1995) and energy minimized. The effect of the mutant was analysed using HOPE (Venselaar et al., 2010) and I-mutant (Capriotti et al., 2006). The I-mutant method allows to predict stability of the protein due to mutation. The docking studies for wild and mutant TMPRSS2 with S-protein and ACE2 were carried out using HADDOCK (Dominguez et al., 2003).

### Network analysis on SARS-CoV-2 with Human Proteins

We downloaded the SARS-CoV-2 genome (Accession number: MT121215) from the NCBI database. In order to find the Host-Pathogen Interactions (HPIs), the SARS-CoV-2 protein sequences were subjected to Host-Pathogen Interaction Database (HPIDB 3.0) (Ammari et al., 2016; Kumar and Nanduri, 2010). In addition to other host genes, we added two more proteins (TMPRSS2 and SLC6A19) which were found to have an important role in the mechanism of viral entry (Hoffmann et al., 2020). The protein-protein interaction (PPI) and transcription factor regulation of human proteins were retrieved from GeneMANIA (Warde-Farley et al., 2010) and literature (Barros et al., 2012; David et al., 2010; Maitland et al., 2011; Tumer et al., 2013; Yu et al., 2010; Zhang et al., 2009), respectively. The Host Pathogen Interaction Network (HPIN) was visualized using Cytoscape (Shannon et al., 2003) which includes the information collected from HPIDB, PPIs and TFs. Modules were defined as the set of statistics and functionally significant interacting genes (Reichardt and Bornholdt, 2006) which was constructed by using MCODE (Bader and Hogue, 2003). Further, Hubs were identified using Network analyzer (Assenov et al., 2008), the plugin of Cytoscape v3.2.1. We also studied the hub proteins' association with diseases, using DisGeNET (Menche et al., 2015).

The topological properties of the network and their functional significance was studied within the formalism of network theory which could predict important organizational and regulatory candidates in the network. We used network theoretical concepts, namely probability of degree distribution, clustering coefficient and average neighborhood connectivity, to characterize the structural and organizational features of the network (Details in supplementary Information). Removal of high degree nodes (hubs) in biological networks may cause lethality to the corresponding organism; the phenomenon has been referred to as the centrality–lethality rule (Jeong et al., 2001), verified by various genomic investigations. This idea of understanding enabled us to determine the topological features of the network architecture, important functionally related modules and hubs (Malik et al., 2019; Nafis et al., 2016). The topological properties were calculated after each subsequent removal, using the Network Analyzer plugin in Cytoscape version 3.7.1. The calculated topological properties of knock-out experiment were compared with those of the main network and the change in the properties helped in understanding the role of leading hubs in the network.

## Results and Discussion

SARS-CoV-2 uses ACE2 for entry, and its S-protein priming by the serine protease TMPRSS2 is a key factor. Despite the availability of other proteases, like cathepsin B and L, in the host cells, yet only TMPRSS2 activity is essential for pathogenesis and spread (Hoffmann et al., 2020).

### Dynamics and binding analysis of ACE2 with SARS CoV-2 RBD in dimeric and monomeric state

Mutagenesis experiments have reported the importance of Arginine and Lysine residues in ACE2, positioned between 697 to 716 as an important recognition site for TMPRSS2 mediated cleavage **(Figure S1a)** of ACE2. Mapping these residues on the recently resolved structure (61M7.pdb) suggests that this region lies in symmetric dimerization interface between ACE2 homodimer; and is also masked from outside by two BoAT1 in an overall 1:2:1 hetero-tetramer of B0AT1 and ACE2 **(Figure S1a)** (Yan and Zhang, 2020). In a recently reported study, the overexpression of *ACE2* has been reported to have an overall protective role in viral infection among patients(Vaduganathan et al., 2020). Increased density of ACE2 on the cell surface has been observed to protect from lung injury (Imai et al., 2005); also observed in many other instances(Kuster et al., 2020). We propose that an increased expression of ACE2 due to various factors, including naturally occurring variations influencing expression of the gene, may increase collisions and hence result in homo-dimerization of this protein. Binding of B0AT1 to ACE2 is not dependent on ACE2 homodimerization, but its presence along with dimerised ACE2 effectively shields the latter from interaction with TMPRSS2, with a possible prevention or reduction in its efficiency to cleave ACE2, a hypothesis based on visualization of the available experimentally resolved structure 61M7.PDB (Yan and Zhang, 2020). This observation is significant since the cleaved ACE2 interacts more efficiently with the SARS CoV-2 RBD domain.

Additionally, we also explored if the protease domains (PD) of each of the homodimerized ACE2 could independently bind with whole of the Spike Glycoprotein without affecting each other or any steric hindrance (Yan and Zhang, 2020). In the absence of any experimentally resolved structure available in literature with ACE2 homodimer and complete SARS-CoV2 Spike Glycoproteins binding to these, we aligned the RBD domains of the 6M17.pdb with the prefusion SARS-CoV2 Spike Glycoprotein in the optimal ACE2 binding conformation (Wrapp et al., 2020; Yan and Zhang, 2020) with “up CTD1” and “open S1 subunit” (Song et al., 2018) (**Figure 1’)**. Both Spike Glycoproteins (6VSB.pdb) aligned well with viral RBD of (6M17.pdb). Further these were also observed without any steric clash with the ACE2 or partner Spike Glycoprotein in modelled structure. Both of these protruded radially outward from ACE2 binding site at PDs but inwards, at anchorage sites on virus membrane, with distance of approximately 27 nanometers (nm) (**Figure 1’)**. However, considering the reported inter spike distance, between 13-15 nm (Neuman et al., 2006) and relative positions of Spike proteins on the viral shell, given almost similar sizes of SARS-CoV (Goldsmith et al., 2004; Neuman et al., 2006) and SARS-CoV-2 (Chen et al., 2020b), these are anticipated to be either parallel or protruding radially outwards from its anchorage site, contrasting to the findings in modelled structure (**Figure 1)**. With this finding of inverse orientation of the Spike protein from virus shell in modelled structure, we propose that in case of ACE2 dimerization, only one of the dimerized ACE2 can bind with the Spike protein due to steric hinderances. Thus, the other unbound ACE2 partner may participate in its physiological role and continue to protect from lung injury.

**Figure 1.**
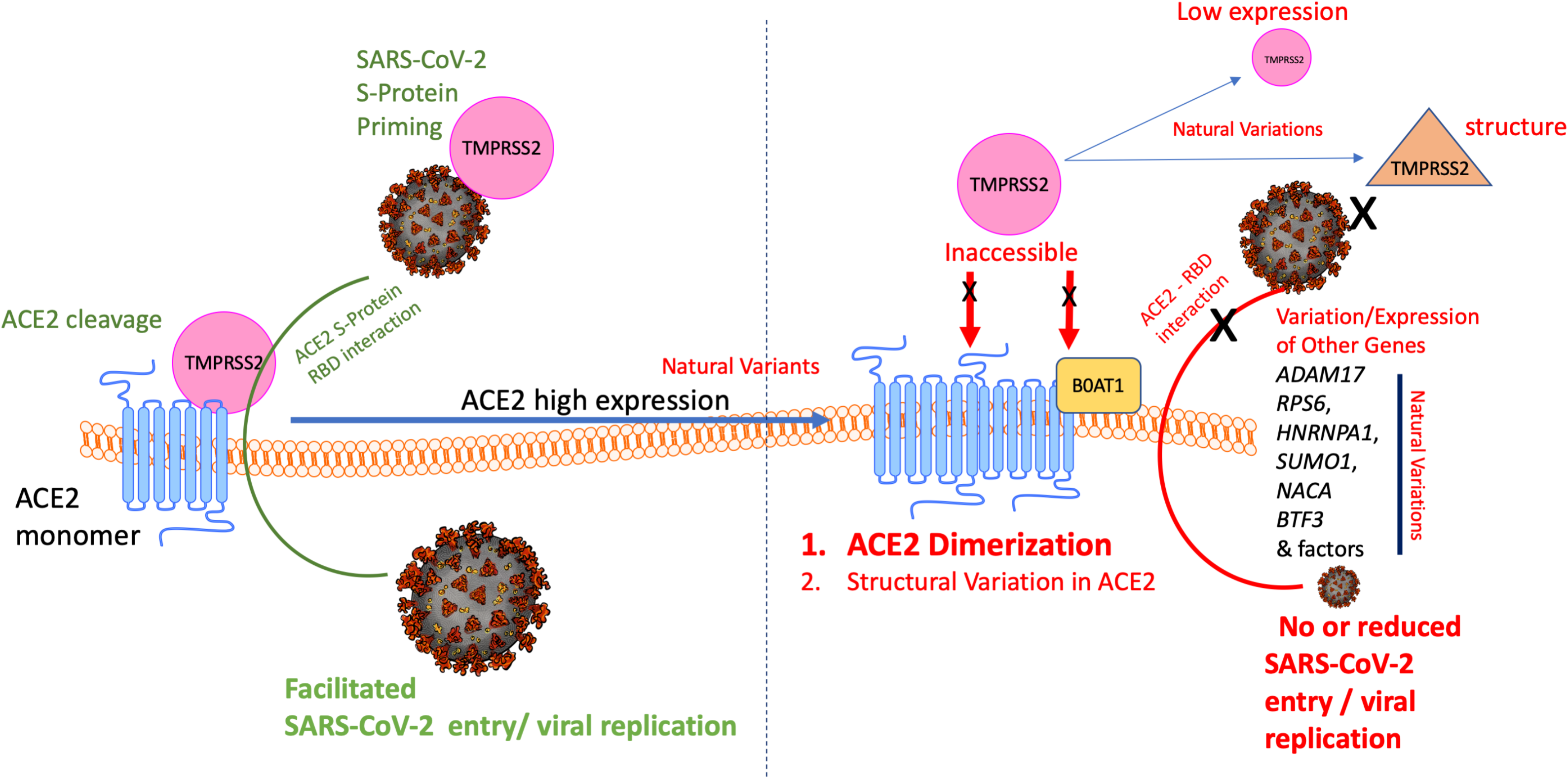
The overall interaction of host proteins and SARS-CoV-2 entry to cell. The figure summarises various factors involved that can influence the entry of Virus in host cell. It depicts brief mechanisms that may arise due to natural occurring variations influencing expression of host genes or structural changes affecting interactions within host proteins or viral proteins resulting in effect on efficiency of virus entry in host cells which could be key factor is providing differential clinical outcomes in different population groups. The figure depicts, higher expression of ACE2 is facilitated by natural variations with different frequencies in different populations and functionally associated with expression of the gene. The higher expression of ACE2 facilitates homo-dimerization resulting in hindrance to TMPRSS2 mediated cleavage of ACE2. Monomeric ACE2 has higher preferential binding with SARS-CoV-2 S-Protein vis-a-vis its dimerized counterpart. It becomes more difficult in presence of B0AT1 that usually does not express in Lungs. Further, natural variations in TMPRSS2, with potential functional role, and their differential frequencies may also result in differential outcomes towards interaction with ACE2 or priming of viral S-protein, a critical step for entry of Virus in host cells. In addition, other potential key host genes like ADAM17, RPS6, HNRNPA1, SUMO1, NACA and BTF3 might have a critical role.

**Figure 1’.**
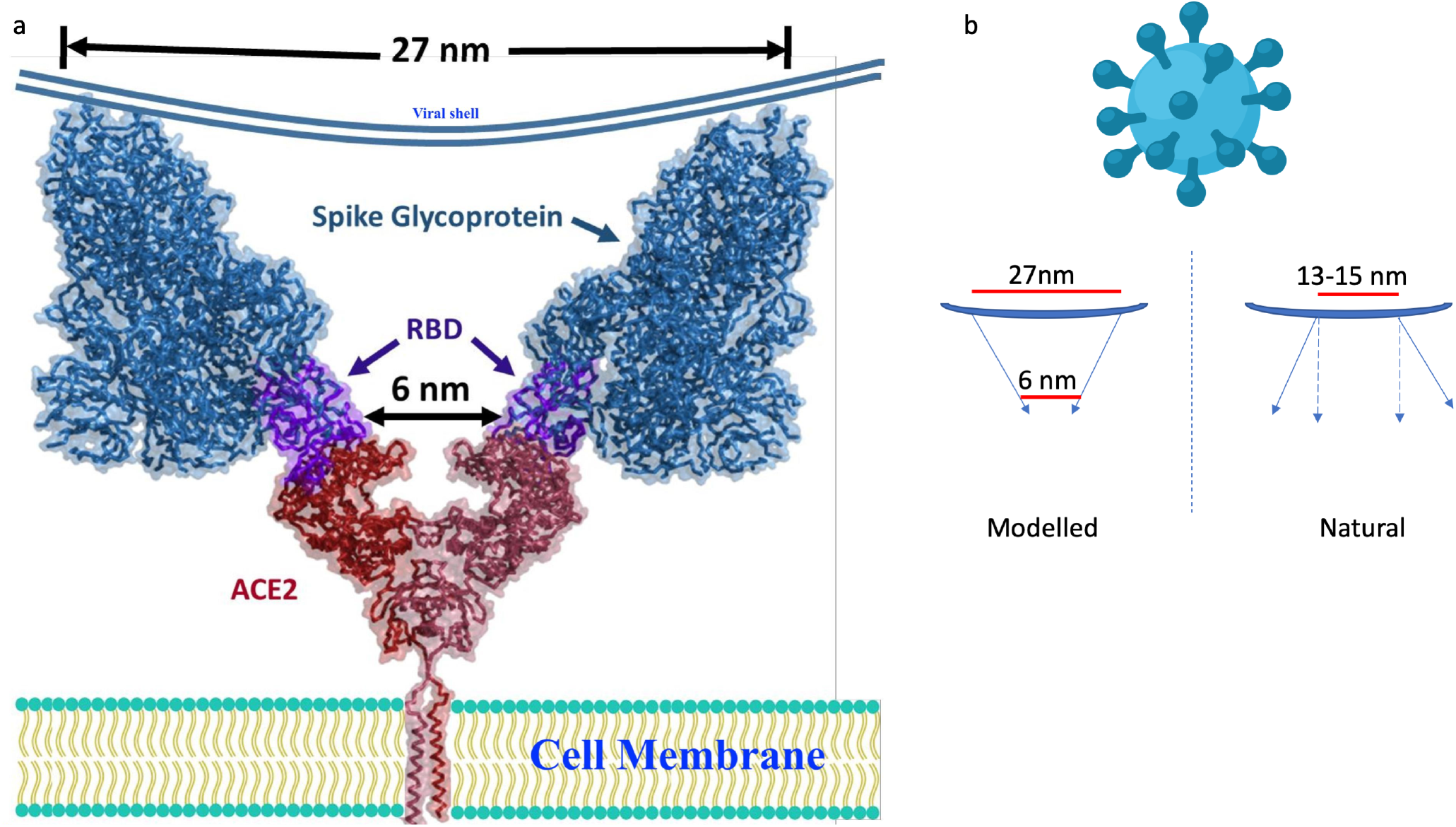
Complex of homo dimer of ACE2 with complete SARS-CoV-2 Spike Glycoprotein. Modelled complex of homo dimer of ACE2 colored as brick red (6M17.pdb) with complete Spike Glycoprotein in open state coloured as sky blue (6VSB.pdb) by aligning the complete Spike Glycoprotein with RBD colored as purple blue from 6M17.pdb. The head of the Spike protein can bind the dimer ACE2, protruding radially outward with RBD domains 6nm apart but inwards, at anchorage sites on virus membrane, with distance of approximately 27 nm.

To understand the mechanistic effect of dimerization, we conceived and simulated two situations: one, where ACE2 dimer is bound to two B0AT1 and two RBD domains of SARS CoV-2; and second, where only monomeric ACE2 is bound to one viral RBD. From the RMSF analysis of the simulated trajectories, we observed a higher order fluctuation in monomeric ACE2 homo-dimerization interface; while the rest of the ACE2 displayed a similar fluctuation profile, including residues interacting with RBD **(Figure S1”)**. PCA analysis and amino acid residue loadings on the PCs (PC1 in this case) reported slightly higher away ward conformational dynamics w.r.t RBD in dimeric ACE2 **(Figure S1”)**. This contrasting essential dynamics in dimeric ACE2 is explained well through the MM-GBSA based free energy of binding calculation between the 10th nanosecond final conformation of ACE2-RBD complex. From both simulations through MM-GBSA method, 1.5 fold strong binding of monomeric ACE2-RBD complex compared to dimeric ACE2-RBD, was observed *(i.e. ΔG*=-75.58 for Dimeric ACE2-RBD vs *ΔG*=-116.98 for Monomeric ACE2-RBD Complex). This observation further explains the protective role of overexpressed ACE2, resulting in its dimerization and reduced affinity for the RBD domain of SARS CoV-2 S1 protein. However, we are also not clear whether the presence of B0AT1 has any allosteric effect on this observation, in addition to already observed masking of ACE2 from TMPRSS2 mediated cleavage. Additionally, another metalloprotease ADAM17 has been reported to compete with the TMPRSS2 and cleave ACE2 in a way that only cleavage by TMPRSS2 was reported to drive the SARS CoV entry inside the cell (Heurich et al., 2014), which is yet to be explored in relation to SARS CoV-2. Recent literature also indicates the potential role of other proteases, like cathepsin B/L that can functionally replace TMPRSS2, which needs to be evaluated extensively. (Sungnak et al., 2020) All these hypotheses warrant further *in-silico* analyses with better computational resources, as well as experimental study designs; and for these we are open to seek collaborations.

Yet, the observations made are of high importance and emphasise on evaluation of expression of the ACE2 and BoAT1 along with TMPRSS2 among patients with varied clinical response to SARS CoV-2 infection, differential outcomes, and correlation with observed mortality. Therefore, a differential expression of these genes among patients displaying varied responses from asymptomatic to acute symptoms is worth an exploration, which may act as biomarkers if proven to predict severity and susceptibility to COVID19.

### Differential Genomics backgrounds derived expression of *ACE2* and *TMPRSS2*

The observations, from our MD analyses **(Figures S1”)** show that higher expression of ACE2 gene may promote ACE2 homo-dimerization, rendering less binding affinity to SARS CoV-2 RBD as well as masking TMPRSS2 cleavage site **(Figure 1).** Based on evidences appearing in literature of eQTLs and ACE2 higher expression in East Asians (Chen et al., 2020a) as well as reported protective role of ACE2 in lung injury (Imai et al., 2005; Kuster et al., 2020; Vaduganathan et al., 2020), we explored more about ACE2 expression and its differential genomic backgrounds in different population groups of the world.

### *ACE2* Expression and functional SNPs related to its expression

Tissue Specific evaluation of ACE2 gene in GTEx portal indicated low level expression of the ACE2 in Lungs (**Figure S2a and S3**). Further, it was observed that not all the transcripts and exons of the gene express in Lungs (**Figure S2b and c**). As expression of *ACE2* is less in Lungs thus, GTEx portal did not return cis-eQTLs for *ACE2* gene in lungs, in all probabilities. However, eQTLs for the gene were found in other tissues **(Figure 2a and b)** in GTEx portal. These eQTLs remained the same across tissue sets and showed similar effect in expression patterns (**Figure 2b)**. Yet to corelate the expression of the gene with genomic variations, regions of the gene were explored for potential regulatory elements and overlapped with common variations through table browser tool of UCSC genome browser with an assumption that the variants having population level effect should be common variants **(Figure S2d and S4).** Variations data was retrieved from 1000 Genomes Phase 3 dataset and filtered for variants with at least 10% frequency of the alternate allele in global population. Overlapping the variants from both the exercises resulted in shortlisting of 2 SNPs rs1978124 and rs2106809, which were observed with differential frequency distribution in different populations of the world **(Figure S5)** at both super-population group and sub-population group level. Interestingly, EAS and EUR sub population groups were observed to show relatively uniform frequency distribution within group. However, in AFR, AMR and SAS sub population groups, intra population group differences were observed to be higher, indicating diversity in the gene pool of population groups. HaploReg annotations indicated that these SNPs are in a region with Enhancer histone marks and DNAase activity. The same were observed through multiple regulatory elements tracks in UCSC Genome browser **(Figure S2d and S4)** including GH0XJ015596 enhancer marked by GeneHancer database. (Fishilevich et al., 2017) Linkage Disequilibrium (LD) values as (r^2^) with other putative proxy functional variants were also explored and mapped with UCSC genome browser. Interestingly, it was observed that SNPs rs1978124 and rs2106809 have a strong LD block of >100kb with various SNPs in absolute LD but upstream of ACE2 gene across population groups **(Figure S6)**. This indicates a strong enhancer activity from the region that may affect higher expression of ACE2 gene in lungs, which may facilitate ACE2 homo-dimerization (**Figure 1**). However, it also requires experimental validation.

**Figure 2.**
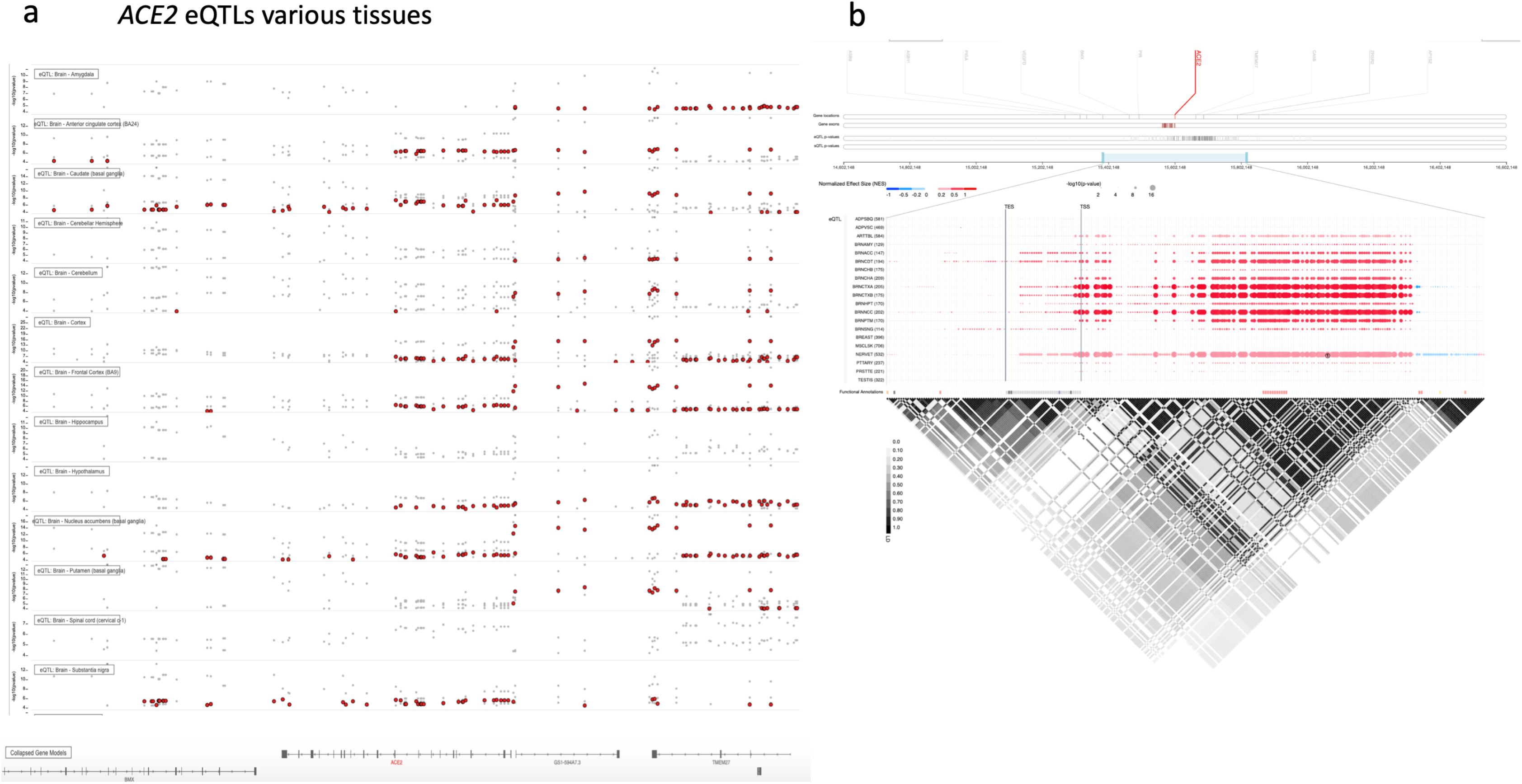
The screenshot from GTEx portal depicting eQTLs of ACE2 Gene. eQTLs in different tissues were plotted through (a) GTEx IGV Browser as well as (b) GTEx Locus Browser. (a) Red dots indicate eQTLs that showed up in the region against the query. Size of the dot in (b) indicate level of significance (as negative p values) whereas colour depicts positive or negative correlation with Normalized effect size (NES) of the eQTL from -1 to 0 to 1. Red color shades represent upregulation whereas blue color shades show downregulation. (b) also depicts Linkage disequilibrium (LD) with value range from 0 to 1 as white to black shades with 1(dark) as absolute LD.

### *TMPRSS2* Expression and functional SNPs in the gene with potential effect

GTEx portal indicated relatively high levels of *TMPRSS2* expression in Lungs (**Figure S2a and S7c**), differential transcription (**Figure S7c**) but no protein expression in lungs (**Figure S7a**). Evaluation of the Protein atlas portal, source of **Figure S7a**, indicated that both antibodies used for detection TMPRSS2 (not shown in Ms) were restricted to target near N terminal of the protein (either cytoplasmic domain or proximal extracellular domain near membrane). Evaluation of the GTEx portal for cis-eQTLs for TMPRSS2 gene in lungs, returned a large number of eQTLs but with a peculiar feature (**Figure 3a and b**). The eQTLs in lungs were different as compared to other tissues and had following features: found to be concentrated in region of the gene with potential alternate transcripts (**Figure a,b and S8**), towards end of the gene in relatively high expressing exons (**Figure 4b)** and coding for the amino acid sequence that has putative functional role in protein as serine protease domain **(Figure 4a)** critical for ACE2 cleavage and SARS CoV-2 S-Protein priming. Variations data was retrieved from 1000 Genomes Phase 3 dataset and filtered for variants with at least 10% frequency of the alternate allele in global population and overlapped with cis-eQTLs data, resulting in 10 SNPs (rs463727, rs55964536, rs4818239, rs734056, rs4290734, rs2276205, rs34783969, rs11702475, rs62217531 and rs383510). Annotation of the SNPs indicated that all the SNPs clustered together and were in strong LD block **(Figure S8 and S9a)**. HaploReg annotations indicated Enhancer histone marks and DNAase activity in the regions overlapping these SNPs. Amongst these, rs4818239 showed a prominent putative functional role (**Figure S9b**) which requires experimental validation. However, the frequency distribution of the variant rs4818239 in different populations groups of 1000G showed an interesting differential pattern (**Figure S9c**). We also explored if there were any alternate functional variations, yet common in populations, by screening TMPRSS2 gene through genomAD browser and filtered for only missense variations. The search returned rs75603675 (NP_001128571.1:p.Gly8Val or G8V) and rs12329760 (NP_001128571.1:p.Val197Met or V197M), also observed with differential frequency distribution in 1000G populations (**Figure S9d and e**). The observations indicate *TMPRSS2* variants may influence interaction with *ACE2* as well as SARS CoV-2 (**Figure 1**) resulting in population specific differential outcomes.

**Figure 3.**
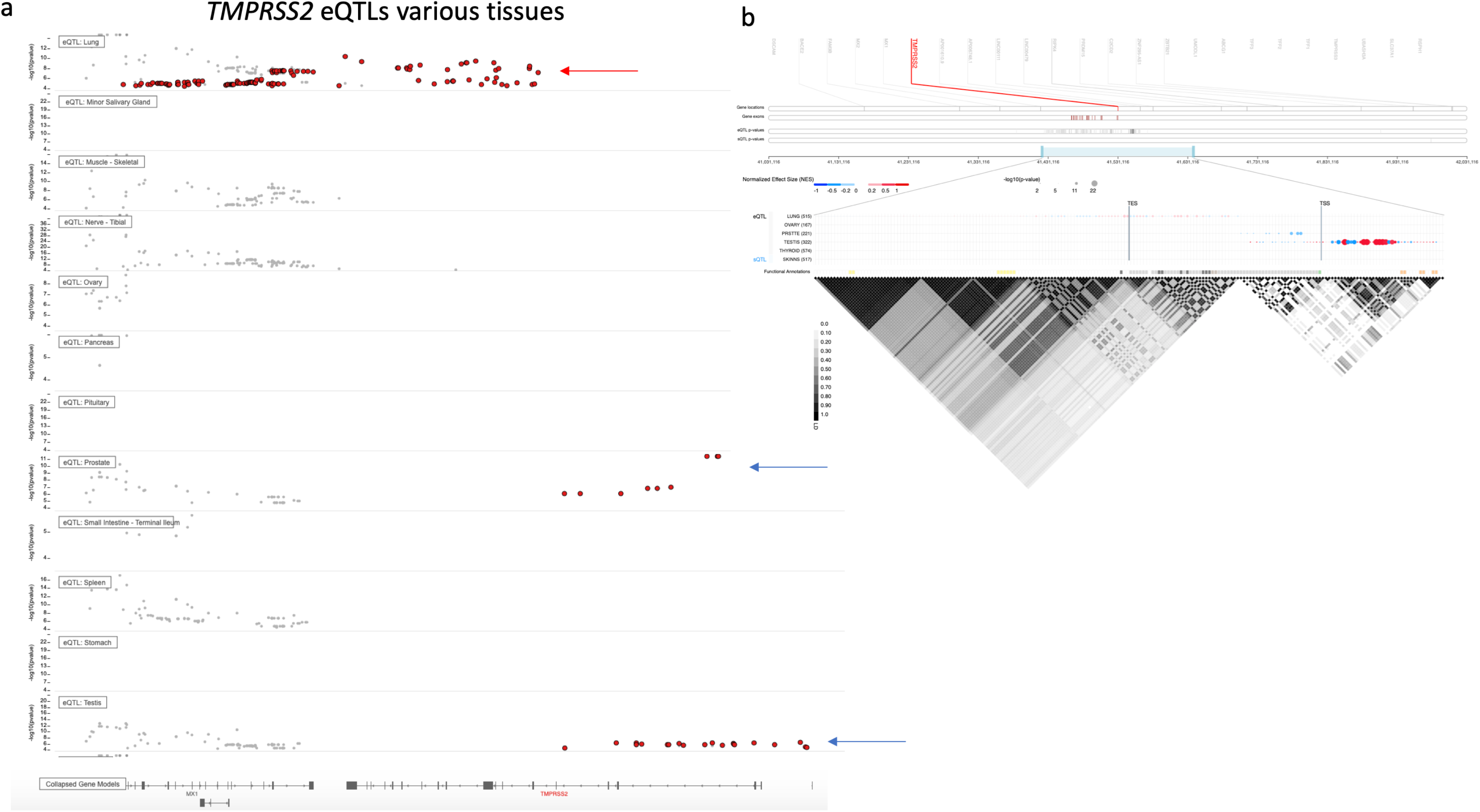
The screenshot from GTEx portal depicting eQTLs of TMPRSS2 gene. eQTLs in different tissues were plotted through (a) GTEx IGV Browser as well as (b) GTEx Locus Browser. (a) Red dots indicate eQTLs that showed up in the region against the query. Size of the dot in (b) indicate level of significance (as negative p values) whereas colour depicts positive or negative correlation with Normalized effect size (NES) of the eQTL from -1 to 0 to 1. Red color shades represent upregulation whereas blue color shades show downregulation. (b) also depicts Linkage disequilibrium (LD) with value range from 0 to 1 as white to black shades with 1(dark) as absolute LD.

It was noted [also indicated by blue arrow in (a)] that where in other tissues eQTLs are mainly towards 5' UTR of the gene, in Lungs the eQTLs are towards end of the gene extending towards 3’UTR.

**Figure 4.**
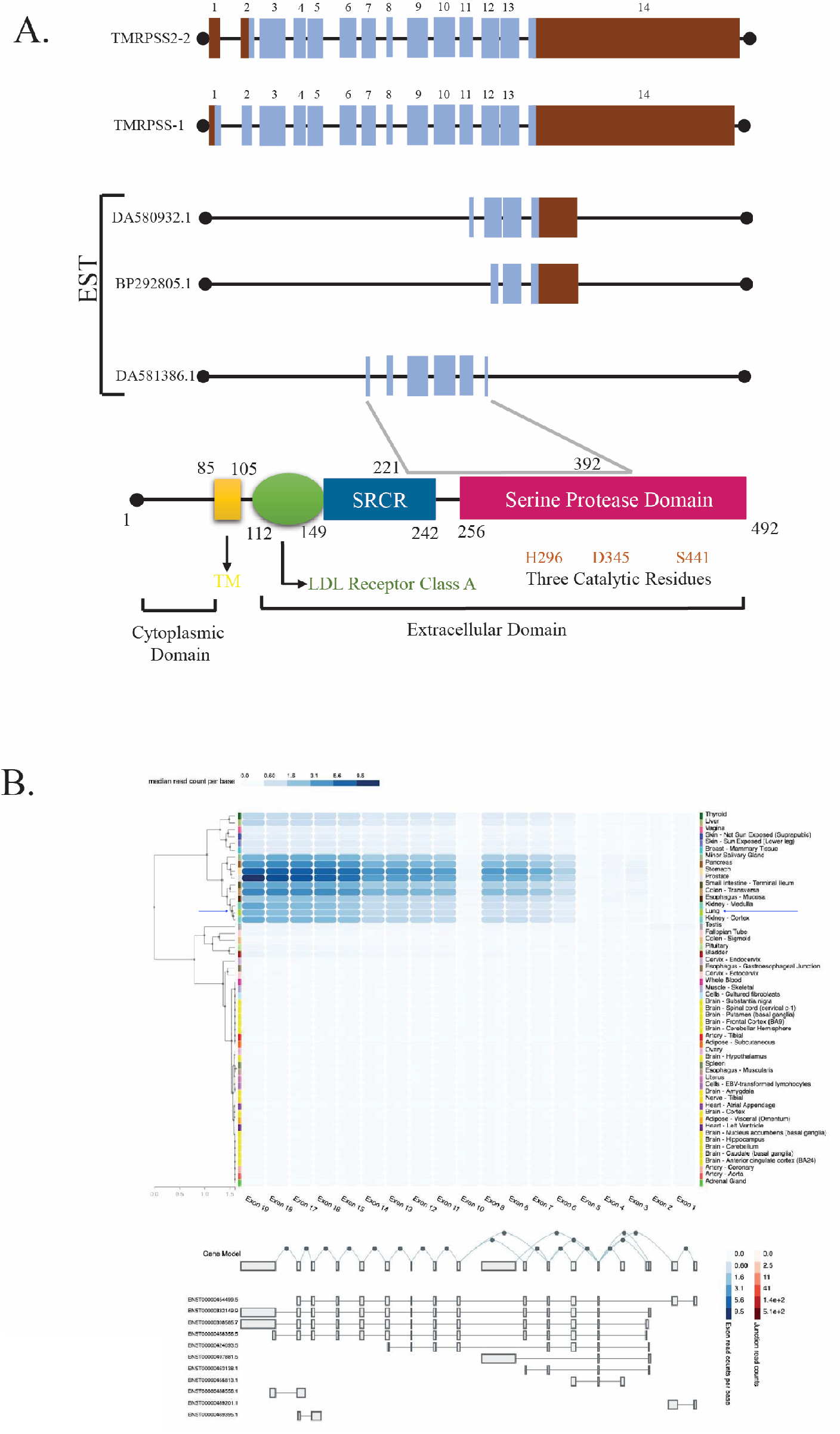
Structure of TMPRSS2 gene, its alternative transcripts and functional domains. (a) TMPRSS2 canonical transcript is constituted of 14 exons and alternate transcripts have been seen. The coded protein is a transmembrane protein with 1-85 amino acids (aa) forming Cytoplasmic Domain, and 112-492 aa constituting Extracellular domain. (b) Expression levels as median read count per base as shades of blue from low to high values indicating exons expression in different tissues in GTEx portal. Gene models of alternate transcripts are also depicted. Interesting to note exons coding for the extracellular domain have higher expression in Lungs (row marked by blue arrow) then other exons.

### The variation in TMPRSS2 could inhibit the ingress of SARS-CoV-2

We further opted to explore the structure and function of TMPRSS2 protein. Pairwise sequence alignment of two isoforms of TMPRSS2 suggested that Isoform-2 lacked 37 residues at N-terminal compared to Isoform-1, which was the longest transcript and coded for 529 amino acids (**Figure S10**). Since the human TMPRSS2 protein structure was not available in the PDB database, we generated a computational protein model. The model was built using Serine protease hepsin (PDB ID: 5CE1) of homo sapiens. The protein 3D structure modelled from 146-491 residues of TMPRSS2 with sequence identity, GMQE and QMEAN of 33.82%, 0.53 and -1.43 values, respectively. It showed that the model was constructed with high confidence and best quality. In a similar manner 3D structure of S-Protein of SARS-CoV-2 was built with sequence identity, GMQE and QMEAN of 99.26%, 0.72 and -2.81 values, respectively. The protein stability analysis showed that rs12329760 (p.Val197Met or V197M) (Isoform-1) variation could decrease the stability of TMPRSS2 protein with ΔΔG value of -1.51 kcal/mol. The HOPE results suggested, V197M variation is located within a domain, SRCR (GO Term: Scavenger Receptor Activity as annotated in UniProt) and introduces an amino acid with different properties, which could disturb this domain and abolish its function. Analyses of another variation rs75603675 (p.Gly8Val or G8V) showed that wild-type residue, glycine, providing flexibility might be necessary for the protein function and an alteration in this position could abolish this function, as the observed torsion angles for this residue were unusual. It could be speculated that only glycine was flexible enough to make these torsion angles, and a change at the location into another residue would force the local backbone into an incorrect conformation, disturbing the local structure. The variant residue was also observed to be more hydrophobic than the wild-type residue. Since sequence similarity search did not find any significant template at the N-terminal of this protein, we could not generate a quality model for isoform-2, which contains G8V variation; hence we carried out sequence-based stability analysis. The results suggested that G8V variation could increase the stability of the TMPRSS2 protein with ΔΔG value of -0.10 kcal/mol. Further, it is known that Arginine and lysine residues within amino acids 697 to 716 are essential for efficient ACE2 cleavage by TMPRSS2 (Heurich et al., 2014) and recent studies have shown that SARS-CoV-2 uses the ACE2 for entry and the serine protease TMPRSS2 for S-protein priming. Based on information from previous studies, we docked the TMPRSS2 p.Val197Met wild-type and variant protein with ACE2 and S-protein of SARS-CoV-2. The docking results suggest that the variant V197M protein could promote the binding to ACE2 and inhibit the binding with S-protein (**Figure S11**). However, these observations need critical revaluation as well as experimental work to understand these interactions better.

### *SLC6A19* (B0AT1) expression naturally in Lungs and other respiratory tract cells, may provide protection

One of the interesting gene, *SLC6A19*, that codes for protein B0AT1, expresses in a very limited number of tissues and is reported to be absent in Lungs (**Figure S2a and S12a,b,c**). Our findings indicated its protective role with competitive hinderance in binding to TMPRSS2. As its expression was observed to be very low in Lungs, we resolved to look for indirect signatures that may have putative role in providing differential susceptibility. We noticed several variations clustering together in a potential enhancer region (**Figure S12d and e**) and with differential frequencies in different populations. eQTL analyses at GTEx portal also showed a huge list of potential SNPs upregulating or down regulating *SLC6A19* expression in Pancreas, liver, and whole blood cells, overlapping with potential Transcription factor binding sites (**Figure S12f)**. We hypothesize a potential chance of some natural occurring variation/s that could induce expression of BoAT1 in respiratory tract thus, providing a protective role in this scenario (**Figure 1**). However, this requires extensive computational exploration as well as experiential validations.

### Host-Pathogen Interaction Modelling

We also believed that there are additional genes and factors which might be playing an additional role in providing differential susceptibility to COVID19. Thus, we carried out a systems biology study to identify the novel key regulators that may influence the Human and SARS CoV-2 interaction. A detailed Host Pathogen Interaction Network (HPIN) was created. The constructed HPIN contained 163 interactions, involving 31 nodes, which included 4 viral proteins, 27 human proteins and 8 Transcription Factors (TF) (**Figure 5a**). From the HPIN, we identified hubs, namely *RPS6, NACA, HNRNPA1, BTF3* and *SUMO1* with 19, 18, 17, 16 & 12 degrees, respectively in the network (**Figure 5a**). This indicated the affinity to attract a large number of low degree nodes towards each hub, which is a strong evidence of controlling the topological properties of the network by these few hubs (Good et al., 2011). Interestingly, out of five significant hubs, four hubs *(RPS6, NACA, HNRNPA1* and *BTF3)* present in module (Nodes: 16, Edges: 118, Score: 15.73) and one hub *SUMO1* was present at motif level which is considered one of the most important regulating motif of biological network at a fundamental level (**Figure 5b**). From our prediction; we found, four viral proteins (*S*, *N*, *ORF1a* & *ORF1ab)* target five host protein groups (*ACE2*, *SUMO1, HNRNPA1, RPS6* & *ATP6V1G1). SUMO1* & *HNRNPA1* are targeted by same viral proteins (N). The hub protein, *RPS6* (highest degree) directly interacted with one of the important protein *TMPRSS2* which further propagated the signal to *ACE2 & SLC6A19*. The Transcription factor HIF1A and BCL6, STAT5, YBX1 inhibits *ACE2* and *SUMO1;* whereas MYC, AR and HNF4a, HNF1a activates *HNRNPA1*, *TMPRSS2* and *SLC6A19* respectively. From the gene diseases association study few diseases were highly associated with these important hub proteins (**Figure S13a**); *HNRNPA1* (36%) followed by *RPS6* (25%), *SUMO1* (21%), *BTF3* (9%) and *NACA* (9%) (**Figure S13b**). Clinically, patients with COVID-19 present with respiratory symptoms, Anoxia, fatigue, heart failure etc could be associated with these hub proteins, mainly *HNRNPA1%#x0026;SUMO1*.

**Figure 5.**
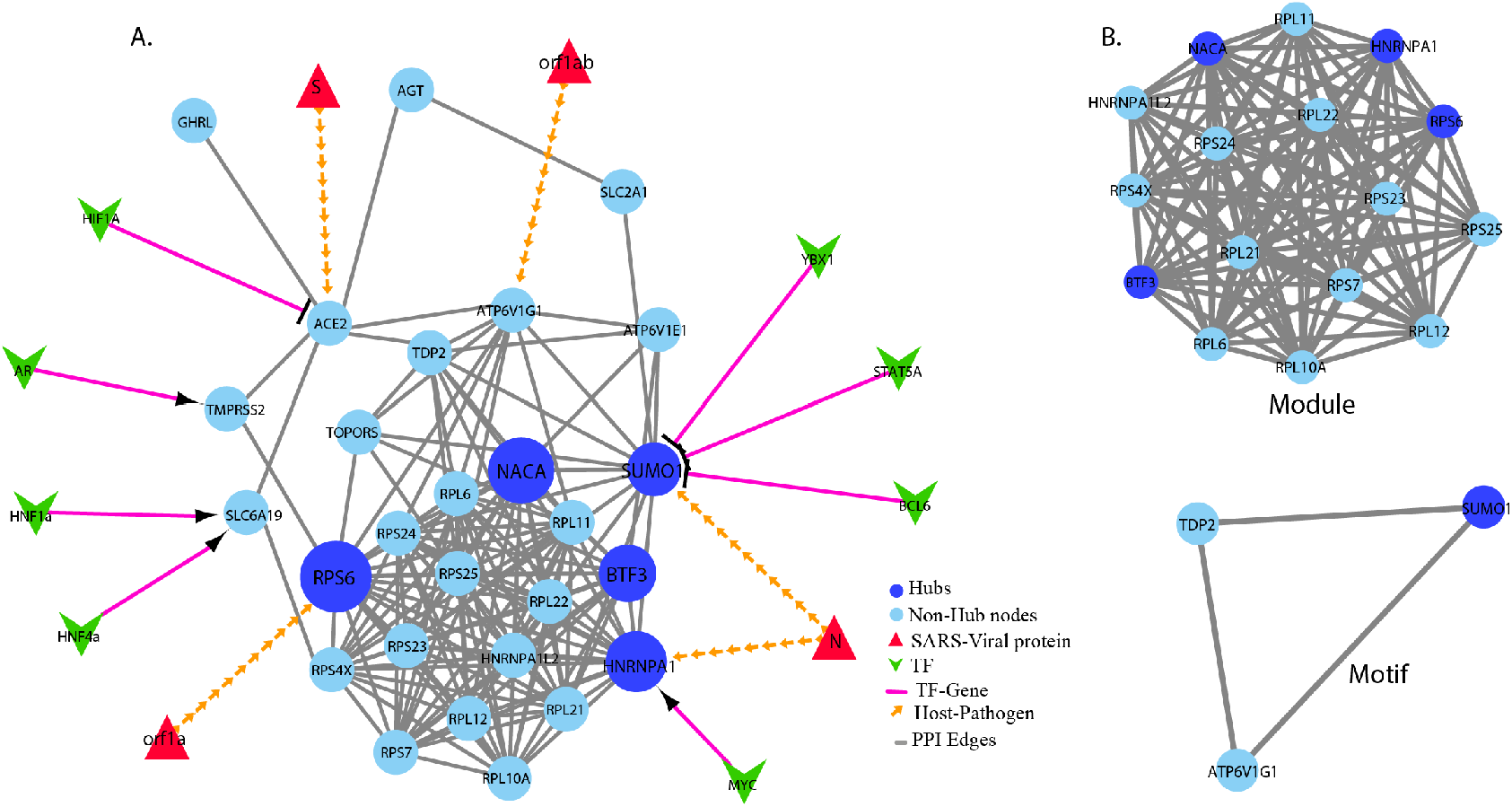
Host-Pathogen Interaction Network and their significant hubs. (a) The network view of HPIN imported from Cytoscape. The Viral proteins, Human proteins and TFs are represented as nodes and edges denote the physical interaction. All the nodes of viral proteins (red), human proteins (blue) and TF (green) are filled triangles, circles and V-shaped respectively. The edges between Virus-human proteins are shown in orange-headed arrows, PPIs in grey lines and TF-human protein in pink arrow-headed (activators) and flat-headed (Inhibitors). The significant existence of sparsely distributed few main hub proteins, namely RPS6, NACA, HNPRNPA1, BTF3 and SUMO1 are colored as dark blue in the network were represented in the order of four enlarged sized circles. (b) The Module and Motif constructed and analyzed using MCODE. All the nodes and edges of module & motif are in blue and grey color, respectively. The significant hubs, present in module & motif highlighted in dark blue color.

By hub removal methodology; we tried to understand the effect of hub removal and calculated the topological properties of the HPIN as a control. The probability of degree distribution (P(k)), Clustering coefficient (C(k)) and Neighbourhood connectivity (C_N_(k)) showed that the HPIN followed a power law scaling behavior. The power law behaviour was also checked and confirmed by using statistical test for power law fitting (p-value ≥ 0.1) (Clauset et al., 2009) in the hub-removal process. The removal of *RPS6, HNRNPA1, SUMO1, NACA* & *BTF3* from the HPIN brings significant variations in the topological properties of the HPIN where, degree distribution (α and β) change significantly (**Figure S14a**). Similarly, the variations in the measurements of the exponents of clustering coefficient and neighbourhood connectivity (λ and μ) also showed significant (Figure S14a). The knockout experiment of hub could able to highlight the local perturbations driven by these hubs and their effect on global network properties. The result suggest removal of *NACA* and *RPS6* hubs could turn down the degree distribution, so it could be crucial for communicating the signals (**Figure S14b**). In case of clustering coefficient, perturbation of *BFT3* shows minimum γ, indicating removal of *BFT3* makes the network less compact, which may lead to the delay in flow of signal. The HPIN perturbation increases in case of *BTF3* hub removal. In Neighbourhood connectivity increase in μ indicates that the information processing in the network becomes faster when *SUMO1* hubs are removed, which means that local perturbations due to removal of hubs are strong enough to cause significant change in global scenario (Canright and Engø-Monsen, 2004) (**Figure S14b**). This indicates hubs are not robust, but it helps in the stability of the network.

Overall network analysis showed *ACE2* is not only the key molecule for entry and survival of SARS-CoV-2 virus, the hub proteins like *RPS6*, *HNRNPA1*, *SUMO1*, *NACA* & *BTF3* might also play a vital role. Analysing these interactions could provide further important understanding for the underlying biological mechanism of SARS-CoV-2 virus infection and identifying putative drug targets.

To conclude (**Figure 1**), higher expression of *ACE2* is facilitated by natural variations with different frequencies in different populations and is functionally associated with expression of the gene. The higher expression of ACE2 facilitates homo-dimerization resulting in hindrance to TMPRSS2 mediated cleavage of ACE2. It becomes more difficult in presence of B0AT1 that usually does not express in Lungs. We also propose that the monomeric ACE2 has higher preferential binding with SARS-CoV-2 S-Protein vis-a-vis its dimerized counterpart. Further, natural variations in *TMPRSS2*, with potential functional role, and their differential frequencies may also result in differential outcomes towards interaction with ACE2 or priming of viral S-protein, a critical step for entry of virus in host cells. In addition, we have identified some other potential key host genes like *ADAM17, RPS6, HNRNPA1, SUMO1, NACA* and *BTF3*, that might have a critical role. With all this background, it is anticipated that in populations like Indian populations, with highly diverse gene pool, a great variation in clinical outcomes is expected and that could be population/region specific, but primarily due to gene pool structure of the region, despite similar exposure levels to SARS CoV-2 and resources. Understanding these population specific differences may help in developing appropriate management strategies.

## Supporting information

ACE2 Homodimerization Affects Binding of SARS-CoV-2 Spike Protein

## Acknowledgment

All the authors acknowledge Prof. RNK Bamezai (Padamshri), former Vice Chancellor Shri Mata Vaishno Devi University, Katra and former Professor and Coordinator, National Centre for Applied Human Genetics, School of Life Sciences, Jawaharlal Nehru University, New Delhi, for his critical suggestions through various virtual discussion rounds, resulting in present study design. Authors also acknowledge Prof. Gyaneshwar Chaubey, Banaras Hindu University, UP, India for inputs, suggestions and assistance with geographical mapping of the allele frequencies from 1000G dataset.

## Author Contributions

SS conceived the concept. IP carried out molecular modelling simulations of ACE2 and related analyses. KP carried out TMPRSS2 modelling and related analyses. SS and ER carried out Human Population data screening and related work. SH and MZM carried out network analyses to identify additional key genes. SS, IP, SH, MZM designed, analysed, executed and wrote their part of manuscript. SS interpreted the results together, ER assisted SS in compilation of figures and SS compiled the overall MS.

## Competing Interests

The authors declare no competing interests associated with MS. For declaration purposes, SS is founder, chief scientific advisor of a startup “Biodroid Innovations Pvt Ltd” and IP is director of “Bioinfores Pvt. Ltd.”.

## Supplementary Information

### Additional information about knock-out experiment in Network analysis

#### Perturbation by leading hub removal analysis

##### Degree Distribution (P(k))

Degree k is the number of interaction a node in the HPIN. *P(k)* is the probability of randomly chosen node to have k interaction with the neighbour. The probability of degree distribution *(P(k))* of the network is calculated by:

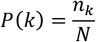

where, *n_k_* is equal to the number of nodes with degree *k* and *N* is equal to the size of the network (Albert and Barabási, 2002; Barabasi and Albert, 1999).

##### Clustering Coefficient (C(k))

Clustering coefficient defines how strongly the nodes in a network tend to cluster together. Clustering coefficient of the *i^th^* node in undirected network can be obtained by:

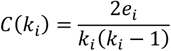

where, *e_i_* is the number of connected pairs of the nearest-neighbour of the *i-th* node and *k_i_* denotes the degree of the respective node (Ravasz and Barabasi, 2003; Ravasz, et al., 2002).

##### **Neighborhood Connectivity** (C_N_(k))

Neighbourhood connectivity of a node is defined as the connectivity of all the neighbours of the node (Maslov and Sneppen, 2002). The average connectivity of the nearest neighbours of a node is given by:

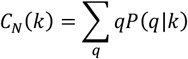

where, *P(q\k)* denotes the conditional probability that a link belonging to a node with connectivity *k* points to a node with connectivity *q*.

For scale free network, *C_N_(k)* is constant while it follows power law *C_N_(k)*= *c* ^−*μ*^ for hierarchical network with μ approximately equal to 0.5 (Pastor-Satorras, et al., 2001). The positive and negative signs in μ indicates the assortivity or disassortivity of network respectively (Barrat, et al., 2004).

## Supplementary figures

**Figure S1.**
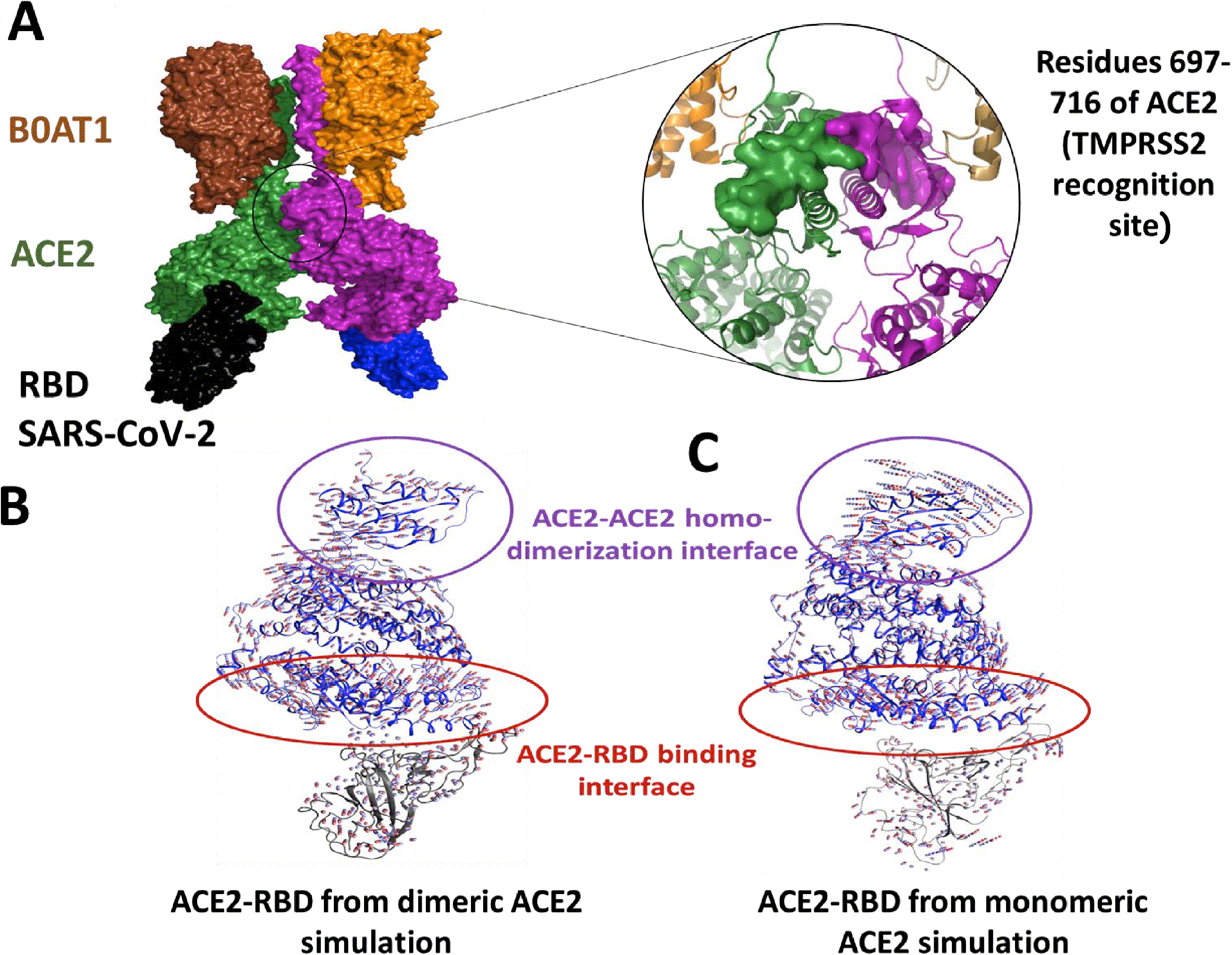
Structural representation and essential dynamics of ACE2. A multi chain complex of dimeric ACE2 is shown (colored as purple and forest green surface representation) bound to B0AT1 (colored as chocolate and orange surface) through its C terminal amino acids and SARS-CoV-2 RBD domain through its N terminal amino acids (shown as black and blue surface), the recognition site of TMPRSS2 is also highlighted as surface against the cartoon background [a]. The cartoon representation along with CPK spheres blue-white-red showing the direction of essential dynamics calculated through PCAs PC1 of ACE2-RBD region from both simulations is shown [b and c].

**Figure S1”.**
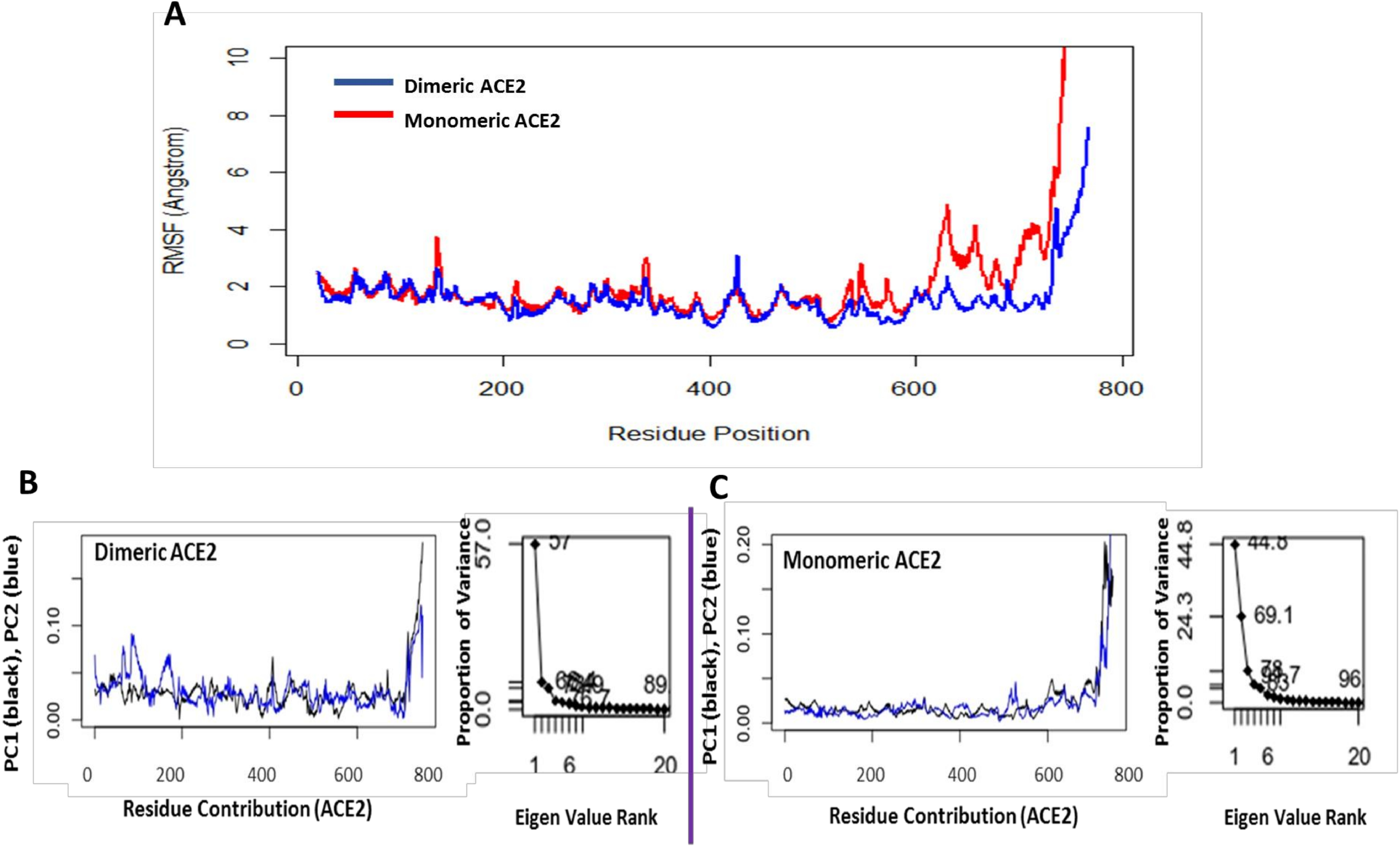
Root mean square fluctuation and contribution of ACE2 residues to PC1 and 2 along with proportion of variance. Root mean square fluctuation (RMSF) plots of ACE2 protein from monomeric ACE2-RBD viral complex and dimeric ACE2(single chain)-RBD complex are shown. The localized fluctuation profile of the amino acids across the protein were similar for both simulation conditions except higher fluctuation in the region that constitutes the ACE2 dimerization interface and following C terminal helix which interacts with B0AT1 (B0AT1 was deleted in the monomeric ACE2-RBD simulation [a]. The contribution of the amino acids to the principal component (PC) 1 and 2 (black and red respectively) along with the scree plot reporting the proportion of variance by each PC are shown [b and c]. It can be observed that the N terminal amino acids consisting of the ACE2-RBD binding site contributed more to the overall essential dynamics in dimeric compared monomeric ACE2 simulated in the present study.

**Figure S2.**
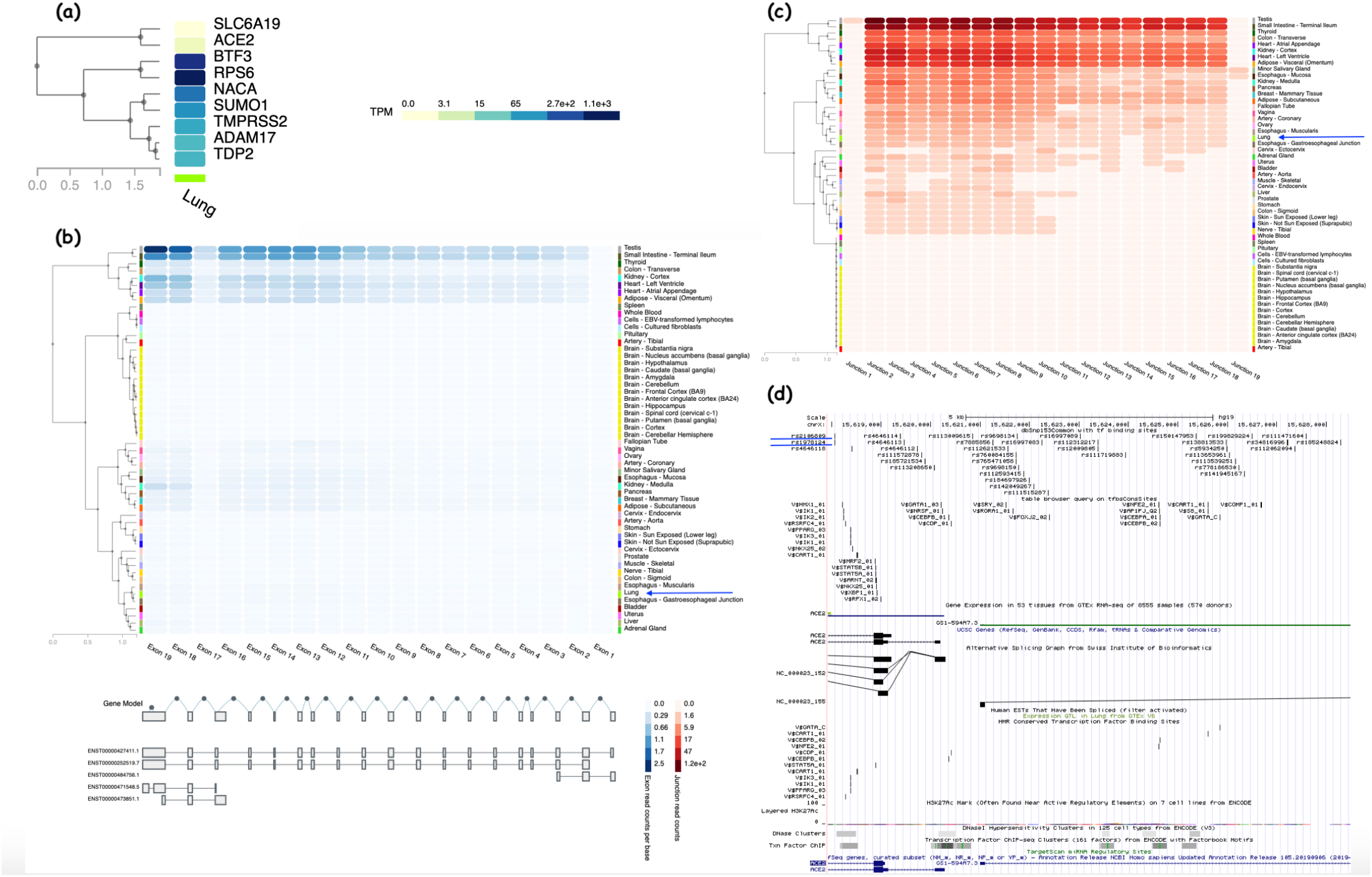
Expression of host genes in Lungs, alternative transcripts of *ACE2* and expression data depiction and regulatory elements and other annotations related to ACE2. (a) Expression of various human genes in Human Lungs from GTEx portal. Normalized Expression Values as Transcripts per million (TPM) are plotted as shades from yellow to deep blue in low to high order. (b) Expression levels as median read count per base as shades of blue from low to high values indicating exons expression in different tissues in GTEx portal. Gene models of alternate transcripts are also depicted. (c) Expression levels as median read count per base as shades of red from low to high values indicating exons junctions expression in different tissues in GTEx portal. (d) UCSC genome browser showing various tracks at 5' Untranslated region of ACE2 gene, common dbSNPs 153 intersecting with conserved transcription factors and conserved transcription factor binding sites are also depicted.

**Figure S3.**
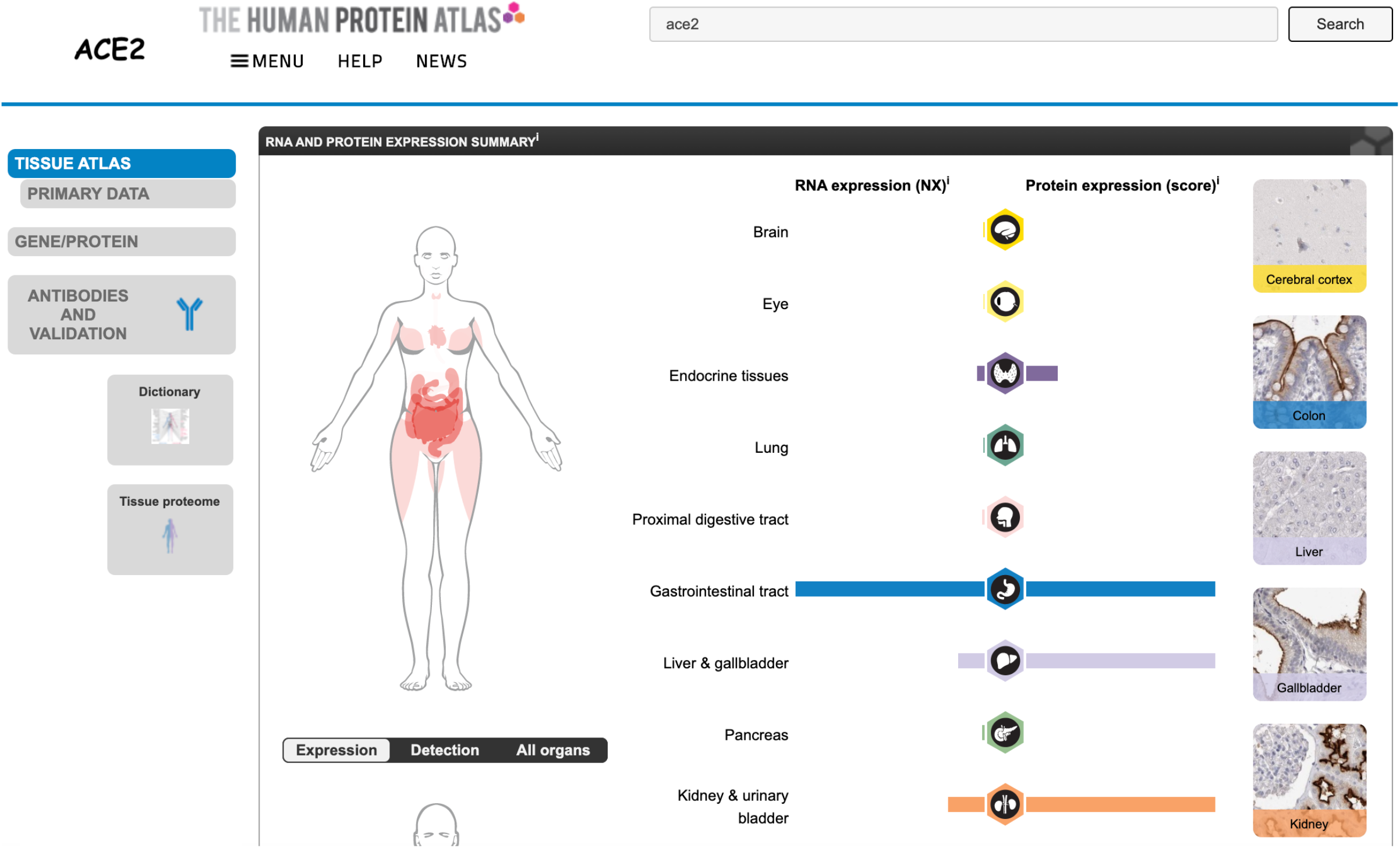
The screenshot from Human Protein Atlas depicting Human ACE2 expression. RNA and Protein expression in different tissues is depicted. In lungs, only RNA expression of ACE2 gene is shown with no ACE2 protein expression.

**Figure S4.**
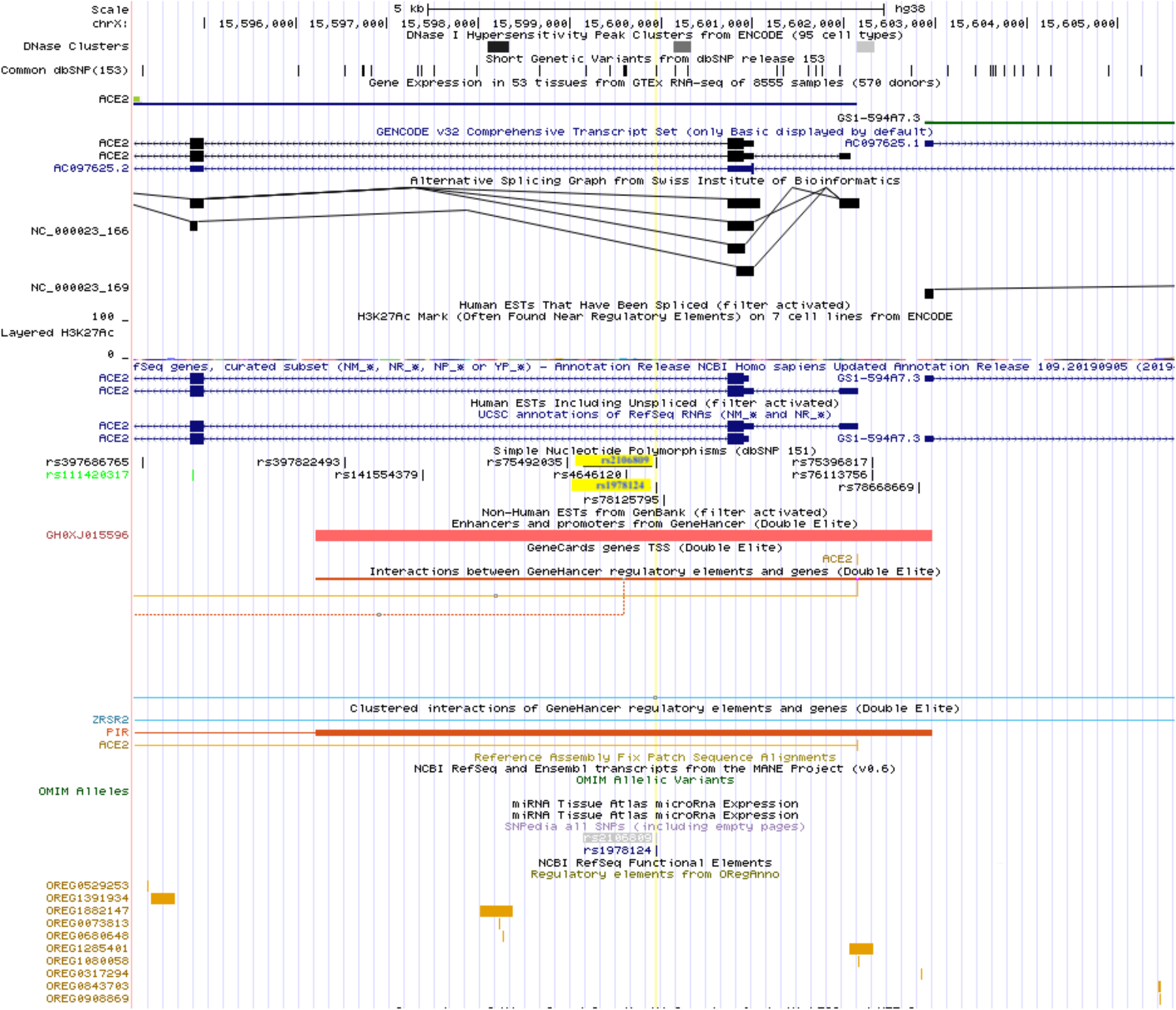
Annotation of *ACE2* depicting various regulatory elements in UCSC Genome browser. UCSC genome browser showing SNPs rs1978124 and rs2106809 in intron 1 of ACE2 gene. Various tracks covering first 2 exons and 5' Untranslated region of ACE2 gene are shown. The ACE2 gene is located on negative strand. In the region GH0XJ015596 enhancer (in red colour) can be located in the track Enhancer and promoter from GeneHancer database.

**Figure S5.**
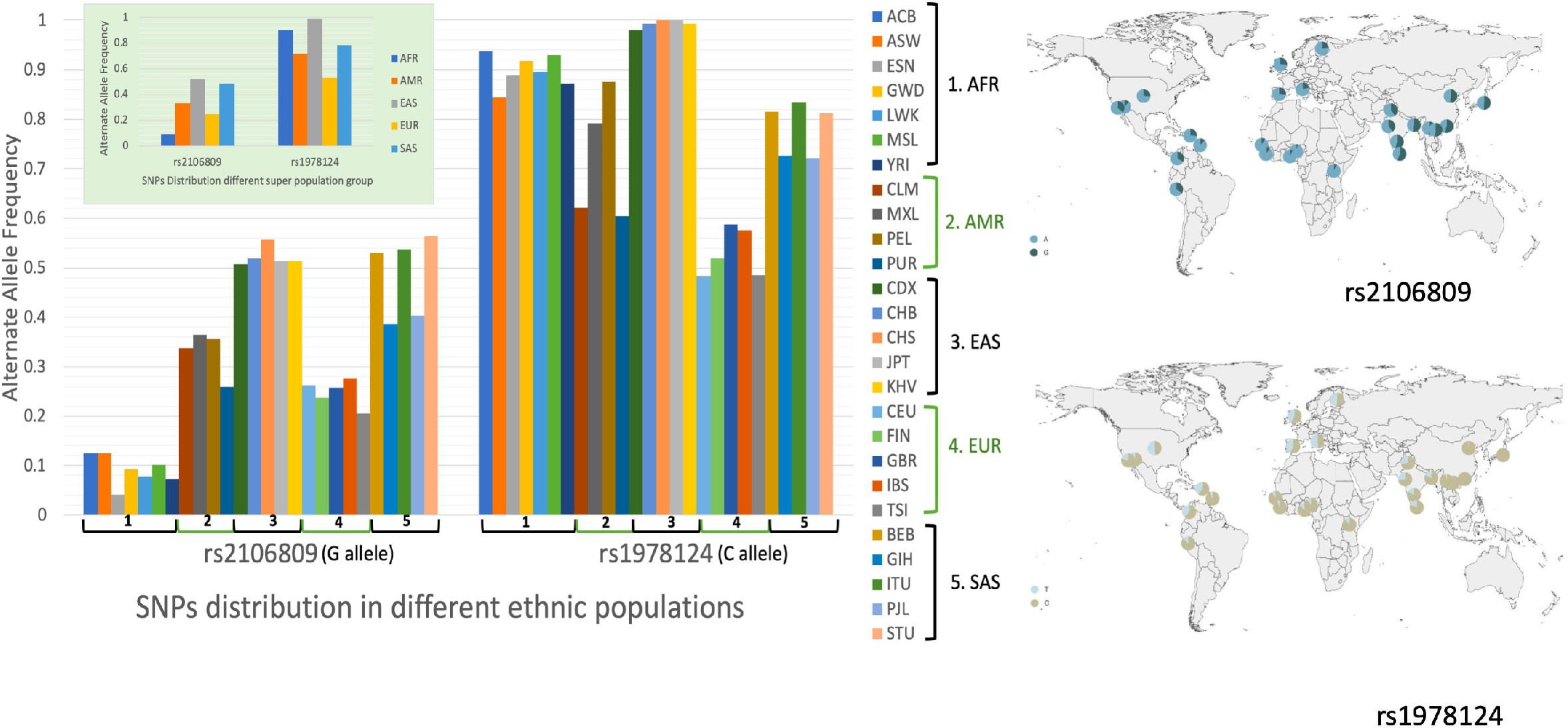
Frequency distribution of SNPs rs1978124 and rs2106809 in 1000G population groups. Frequency distribution of SNPs, rs1978124 and rs2106809 in 1000G populations dataset, also depicted on world map by a web based tool Datawrapper (https://www.datawrapper.de/). Derived allele frequencies in both the SNPs are also plotted for comparison, indicating differential frequency distribution in different population groups. SNPs showed relatively uniform frequency distribution trend in sub populations belonging to same super population groups EUR and EAS. However, differences can be seen amongst the AFR, AMR and SAS sub population groups.

**Figure S6.**
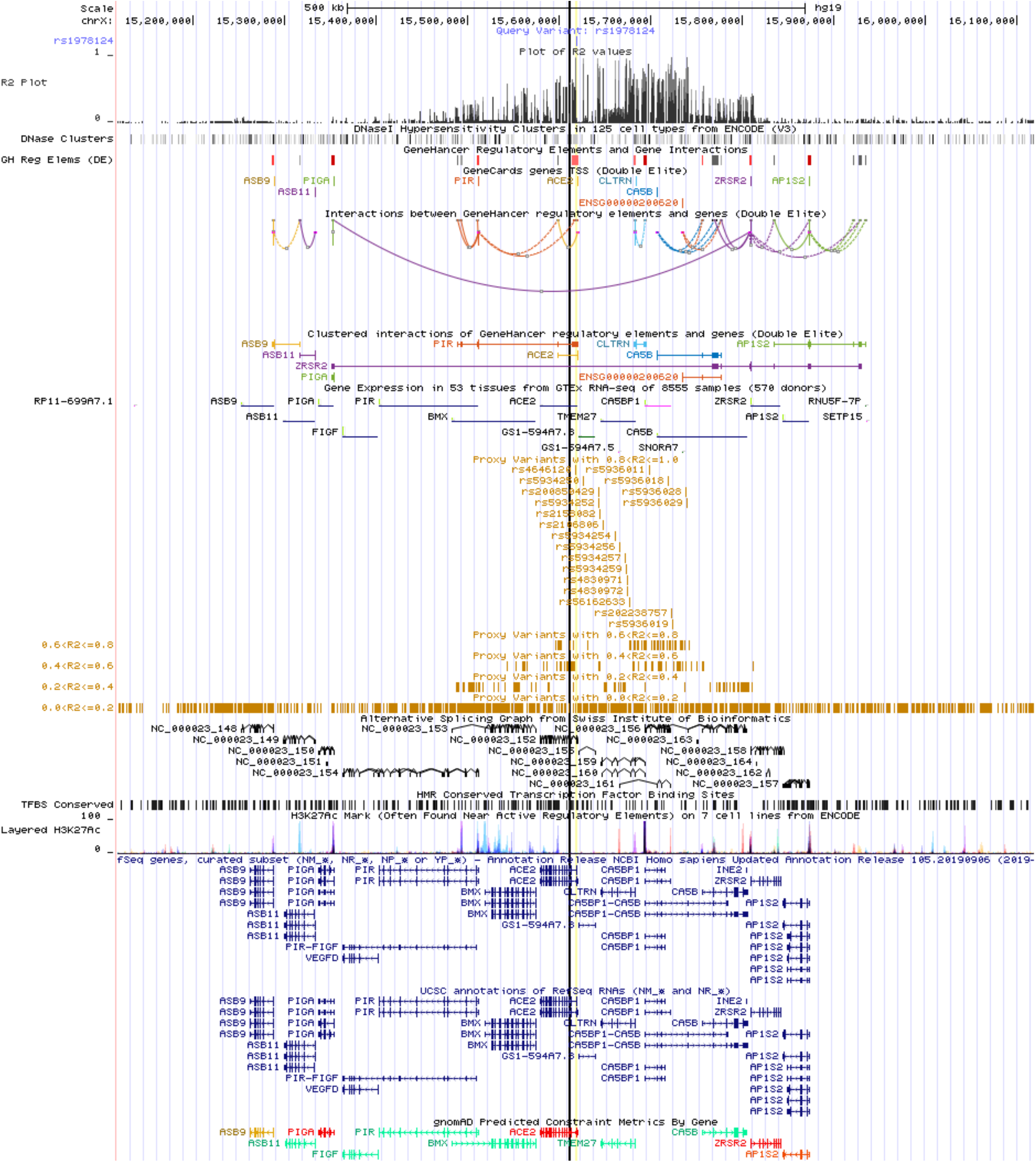
UCSC Genome browser screenshot depicting proxy SNPs and LD structure in the *ACE2* region. UCSC genome browser showing r^2^ plot along with other annotations for SNPs rs1978124 and rs2106809 from the intron 1 of ACE2 gene. GeneHancer regulator elements and interaction between GeneHancer regulatory elements as curved lines can be seen. In addition, it can be seen that all the proxy SNPs (in Brown) are clustered together within potential functional regulatory enhancer element and interaction region (in brown color).

**Figure S7.**
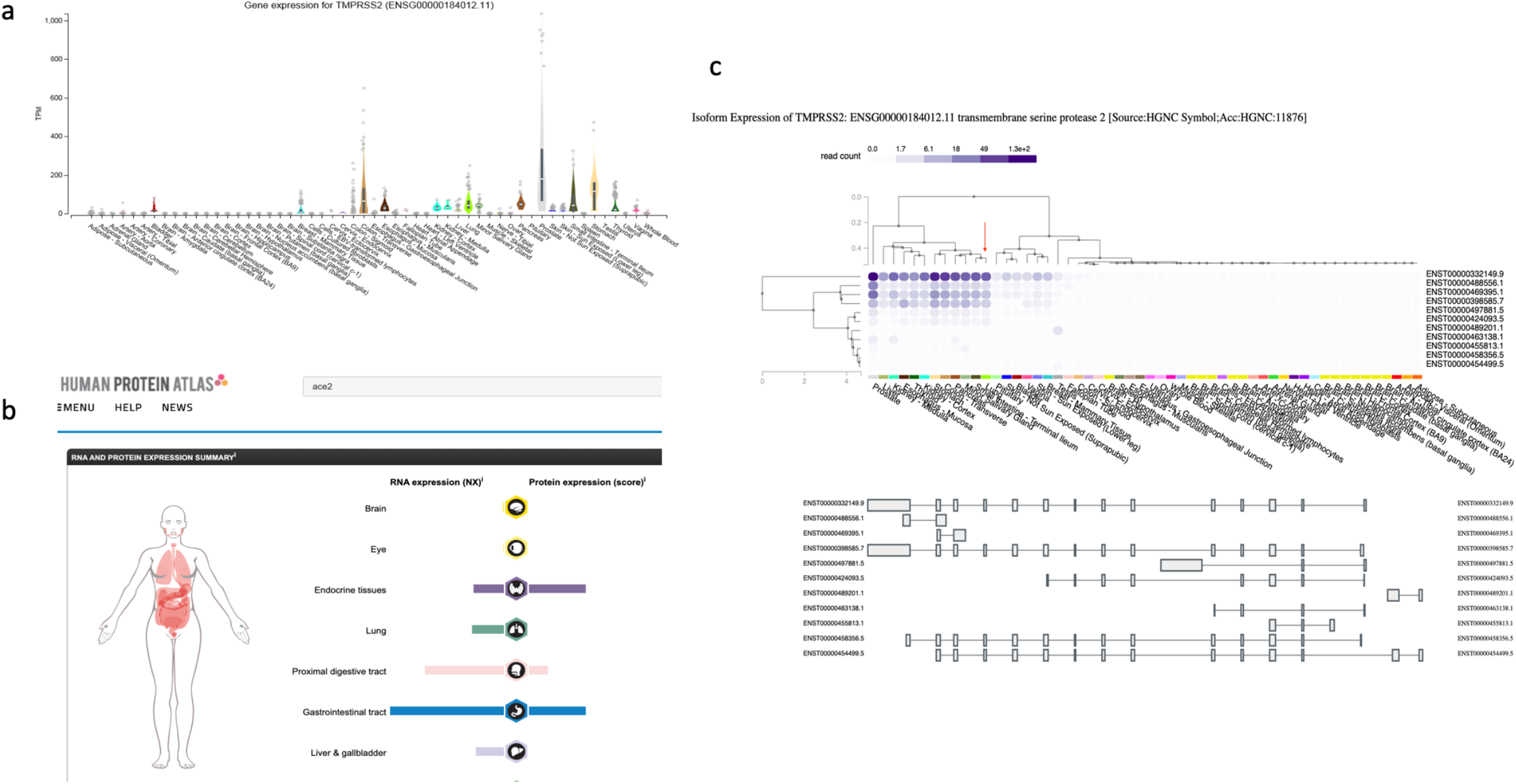
Expression of TMPRSS2 in different tissues. (a) Expression of TMPRSS2 in different tissues as reterived from GTEx portal. Transcripts per million (TPM) are plotted on Y axis and different tissues on X axis. (b) RNA and Protein expression in different tissues is depicted in Human Protein Atlas online portal. In lungs, only RNA expression of TMPRSS2 gene is noted with no protein expression depiction. (c) Expression levels as read count in shades of purple from low to high values indicating expression of different transcripts in different tissues in GTEx portal. Gene models of alternate transcripts are also depicted.

**Figure S8.**
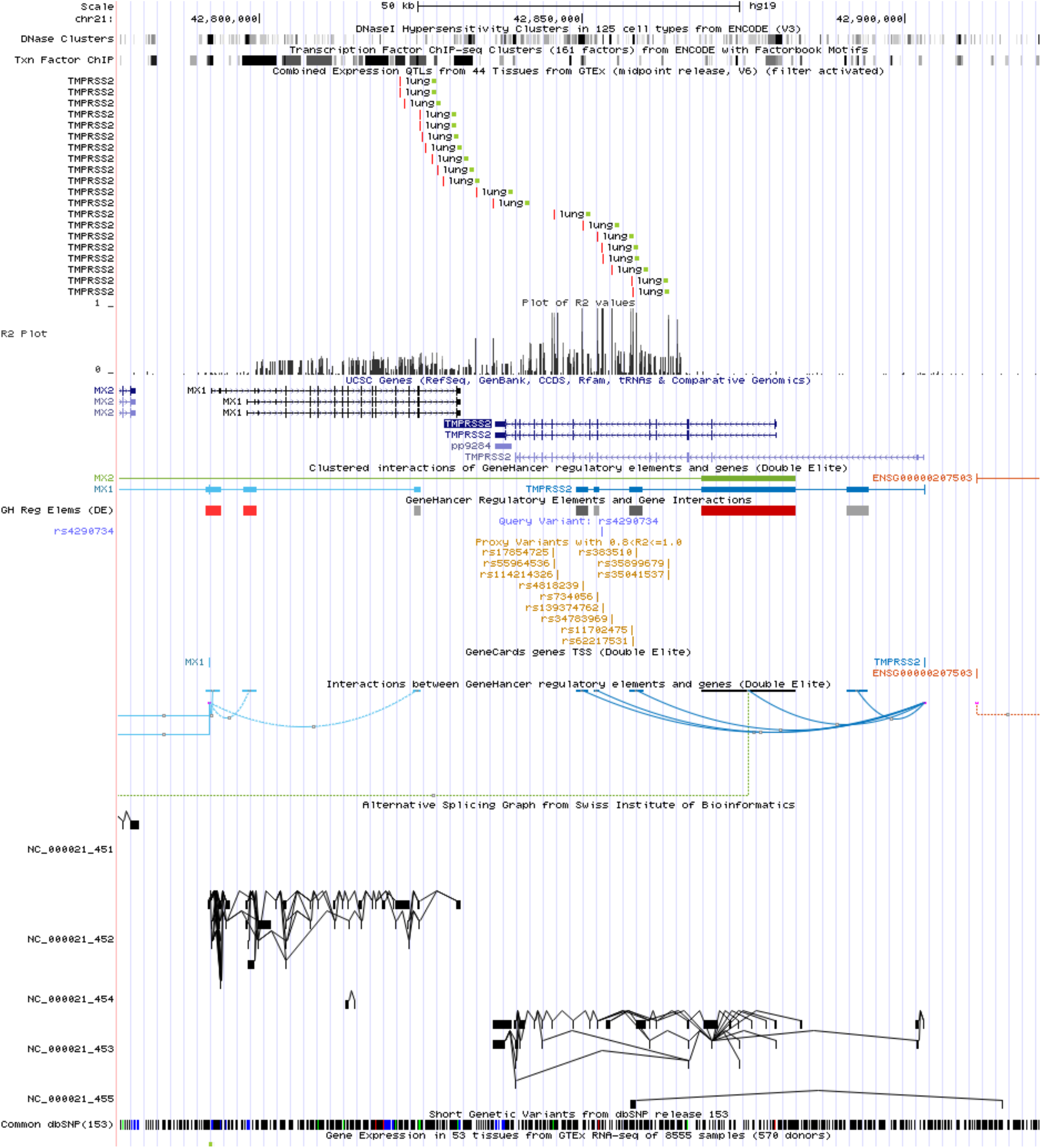
UCSC Genome browser screenshot depicting Lung eQTLs along with SNPs and LD structure in the *TMPRSS2* region. Lung eQTLs that overlapped with the common dbSNPs are shown and along r^2^ values depicting LD in the region is plotted. All the SNPs appeared to cluster towards 3' end of the gene. Other annotations include GeneHancer regulator elements and interaction between GeneHancer regulatory elements as curved lines can be seen. (dark blue color for TMPRSS2 gene). Alternate splicing graph is also indicated.

**Figure S9.**
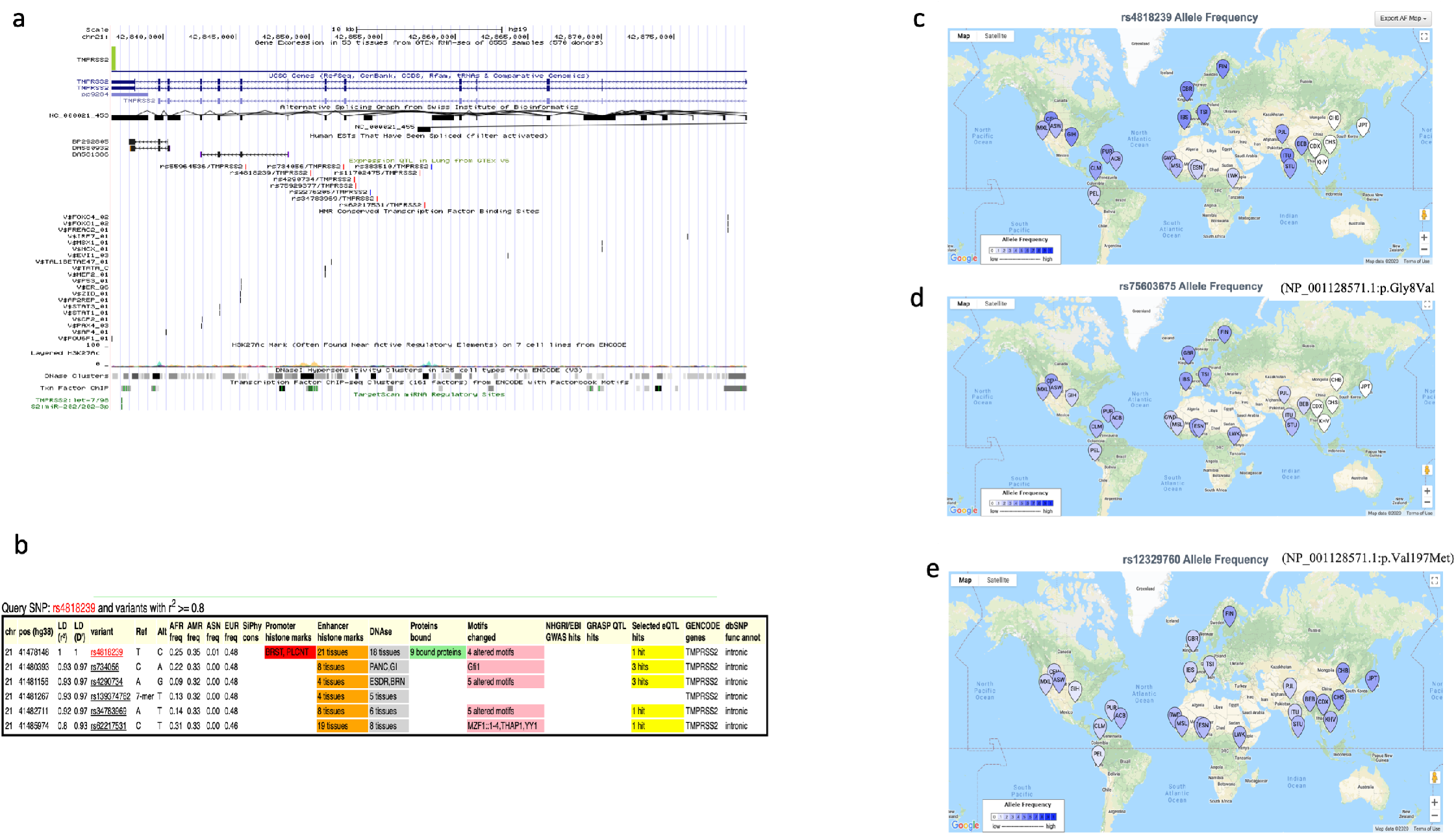
Elucidation of key functional SNPs of *TMPRSS2* and their Frequency distribution in 1000G population groups. (a) TMPRSS2 gene eQTLs in lungs appear to cluster in the gene towards 3' end and potentially are associated with expression of alternative transcripts in lungs. (b) Amongst these common eQTLs, HaploReg annotations indicated rs4818239 as an important SNP. (c) Frequency distribution on world map of rs4818239 (d) rs75603675 (p.Gly8Val) (e) rs12329760 (p.Val197Met) The frequencies of SNPs are based on 1000G populations dataset and depicted on world map by LDpop tool of web based LDlink 4.0.3 suite from National Cancer Institute, USA. Derived allele frequencies from low to high are plotted as shades of white to blue.

**Figure S10.**
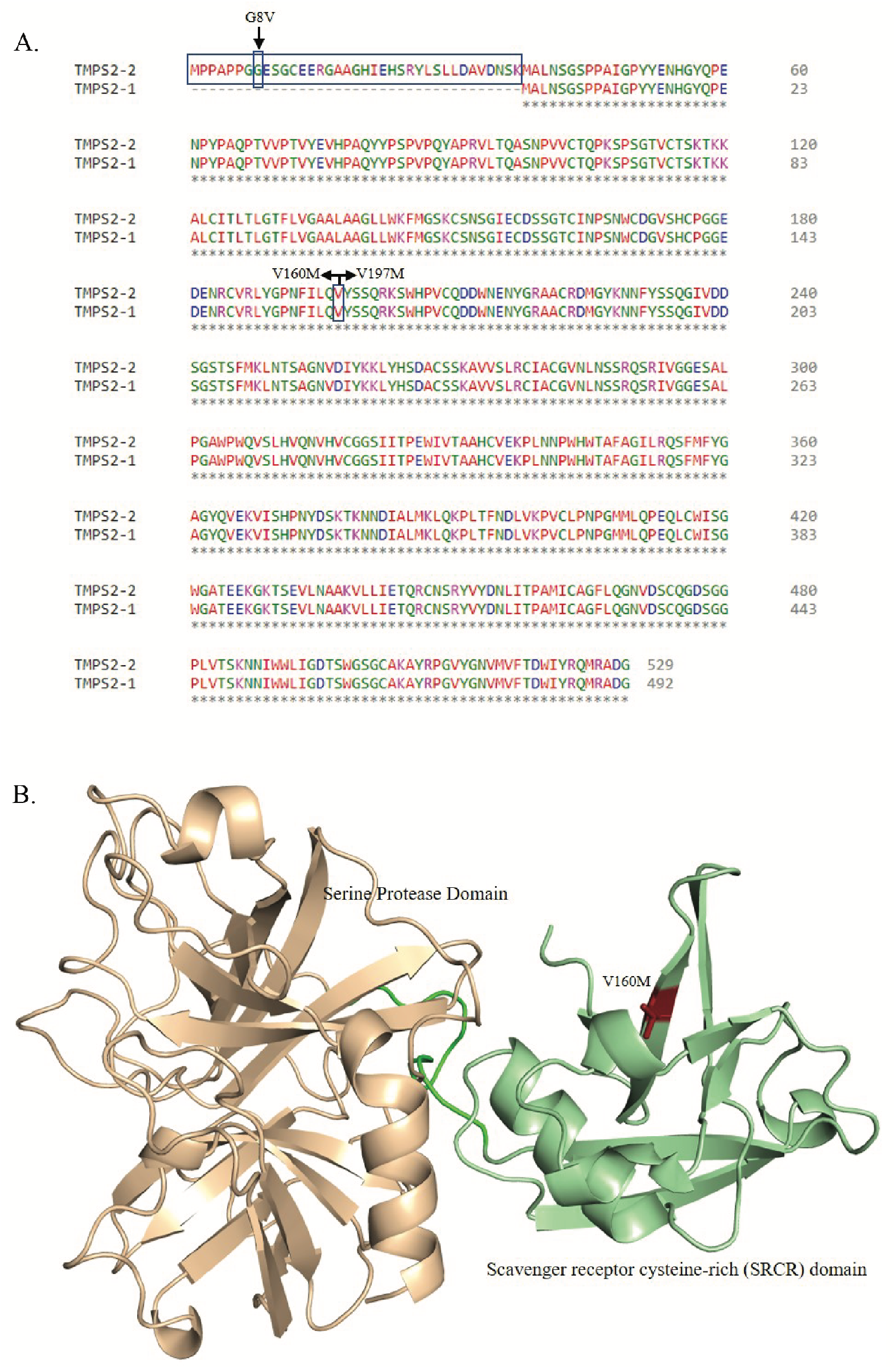
Sequence alignment and structure of TMPRSS2. (a) The pairwise sequence alignment shows the N-terminal difference in two isoforms and highlights two mutation regions. (b) The predicted 3D structure of TMPRSS2 protein. Two domains of TMPRSS2 and mutation residue are highlighted in the carton model with different colors.

**Figure S11.**
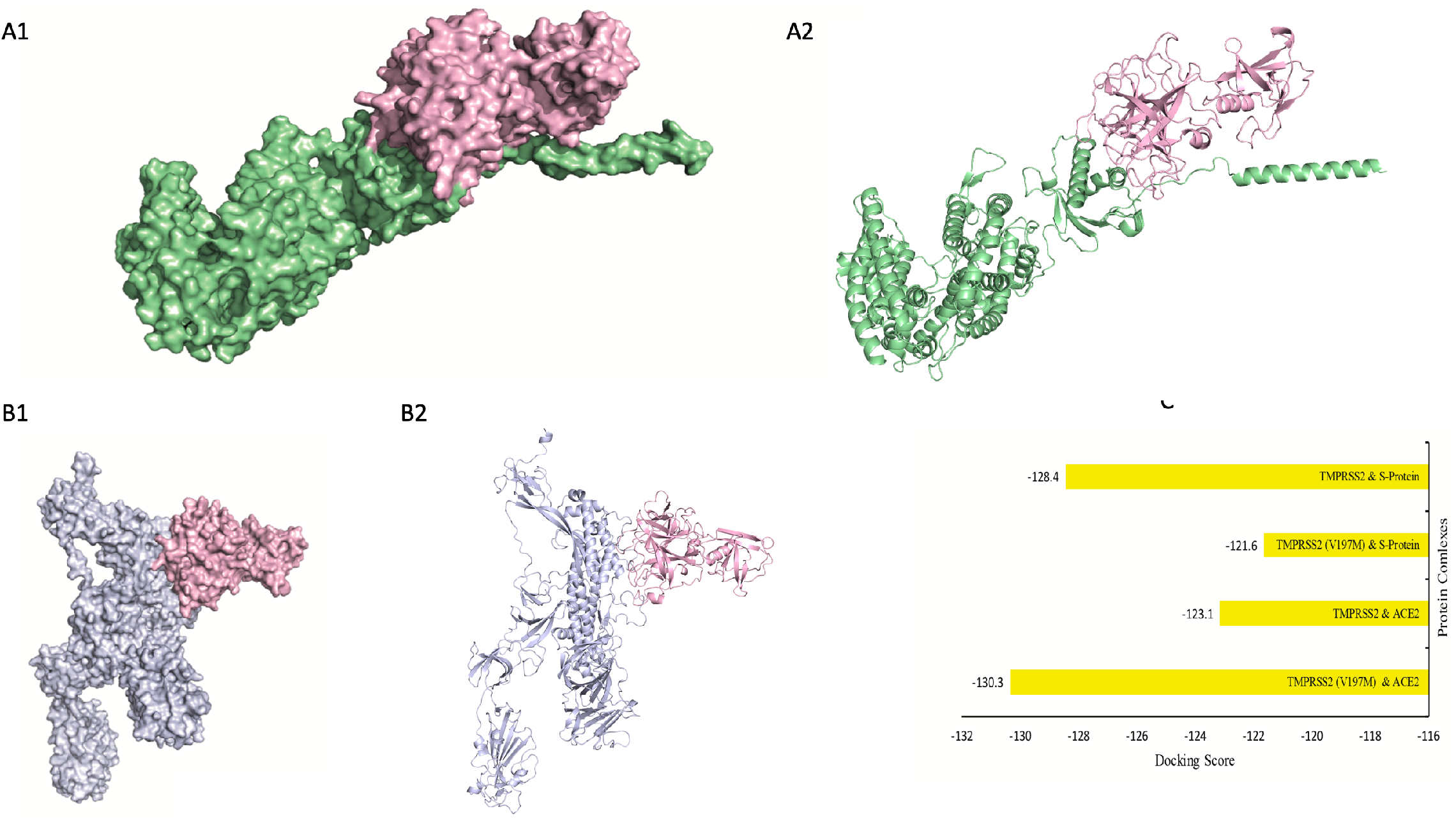
*In silico* protein-protein docking analysis of TMPRSS2. (A1) Surface Model (A2) Carton Model of TMPRSS2 (Pink) human protein interaction with ACE2 (Green) of SARS CoV-2. (B1) Surface Model (B2) Carton Model of Docked complex of SARS CoV-2 S-protein protein (Blue) with TMPRSS2 (Pink). (C) Comparison of docking score between wild-type and mutant TMPRSS2 with ACE2 and SARS CoV-2 S-protein.

**Figure S12.**
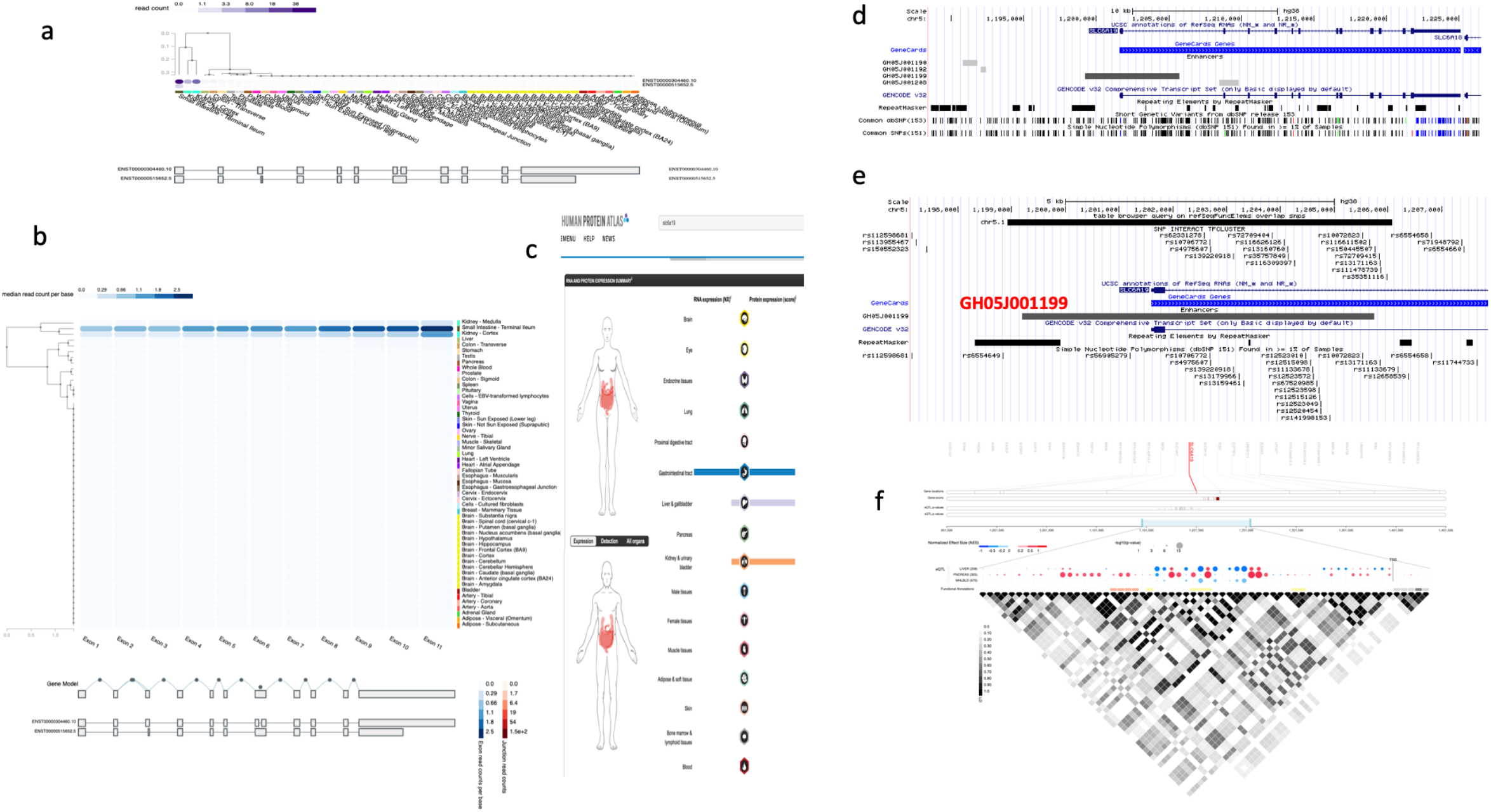
Summarised information about SLC6A19 gene and its variants. (a) Expression of TMPRSS2 in different tissues as retrieved from GTEx portal indicates very restricted expression of the gene. Expression levels as read count in shades of purple from low to high values. Gene models of alternate transcripts are also depicted.(b) Expression levels as median read count per base as shades of blue from low to high values indicating exons expression in different tissues in GTEx portal. Gene models of alternate transcripts are also depicted. (c) SLC6A19 RNA and Protein expression in different tissues is depicted in Human Protein Atlas online portal indicating SLC6A19 has very restricted expression in tissues and in lungs neither its RNA nor Protein expresses. (d) UCSC genome browser screenshot highlights a prominent Enhancer element GH05J001199 at 5'UTR of the gene overlapping exon 1 and extending in intron 1 of the gene. (e) Screenshot from UCSC genome browser depicting SNPs cluster overlapping and intersecting enhancer region GH05J001199. It also depicts common variants (with applied filter on track for SNPs with frequency greater than 20 percent in global populations) in dbSNP version 151. (f) eQTLs in different tissues for *SLC6A19* were plotted through GTEx Locus Browser. Size of the dot indicate level of significance (as negative p values) whereas colour depicts positive or negative correlation with Normalized effect size (NES) of the eQTL from -1 to 0 to 1. Red color shades represent upregulation whereas blue color shades show downregulation. It also depicts Linkage disequilibrium (LD) in region with value range from 0 to 1 as white to black shades with 1(dark) as absolute LD. Significant upregulation of the gene by eQTLs was observed in Pancreas whereas downregulation in Liver and no data on Lungs as it is reported not to be expressing in Lungs at GTEx portal.

**Figure S13.**
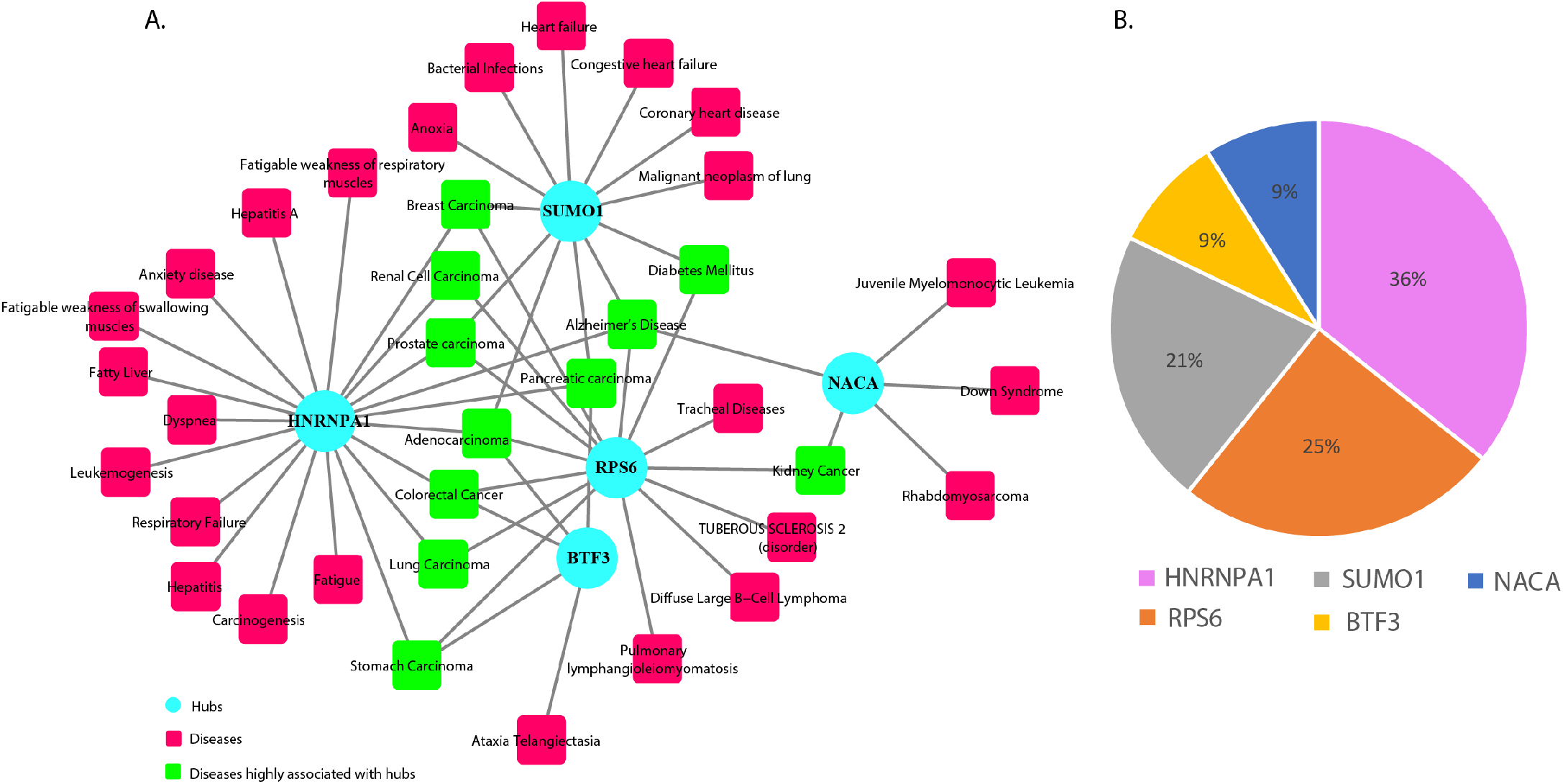
Number of diseases associated with hub proteins. (a) The network view where nodes represent diseases and hub proteins and edges the association between them. (All the nodes of diseases rectangle (red), diseases highly associated with hub proteins rectangle (light green) and hub proteins circle (cyan) are filled color and edges in lines (grey). (b) The pie chart graph showing the percentage distribution of each hub proteins associated with diseases.

**Figure S14.**
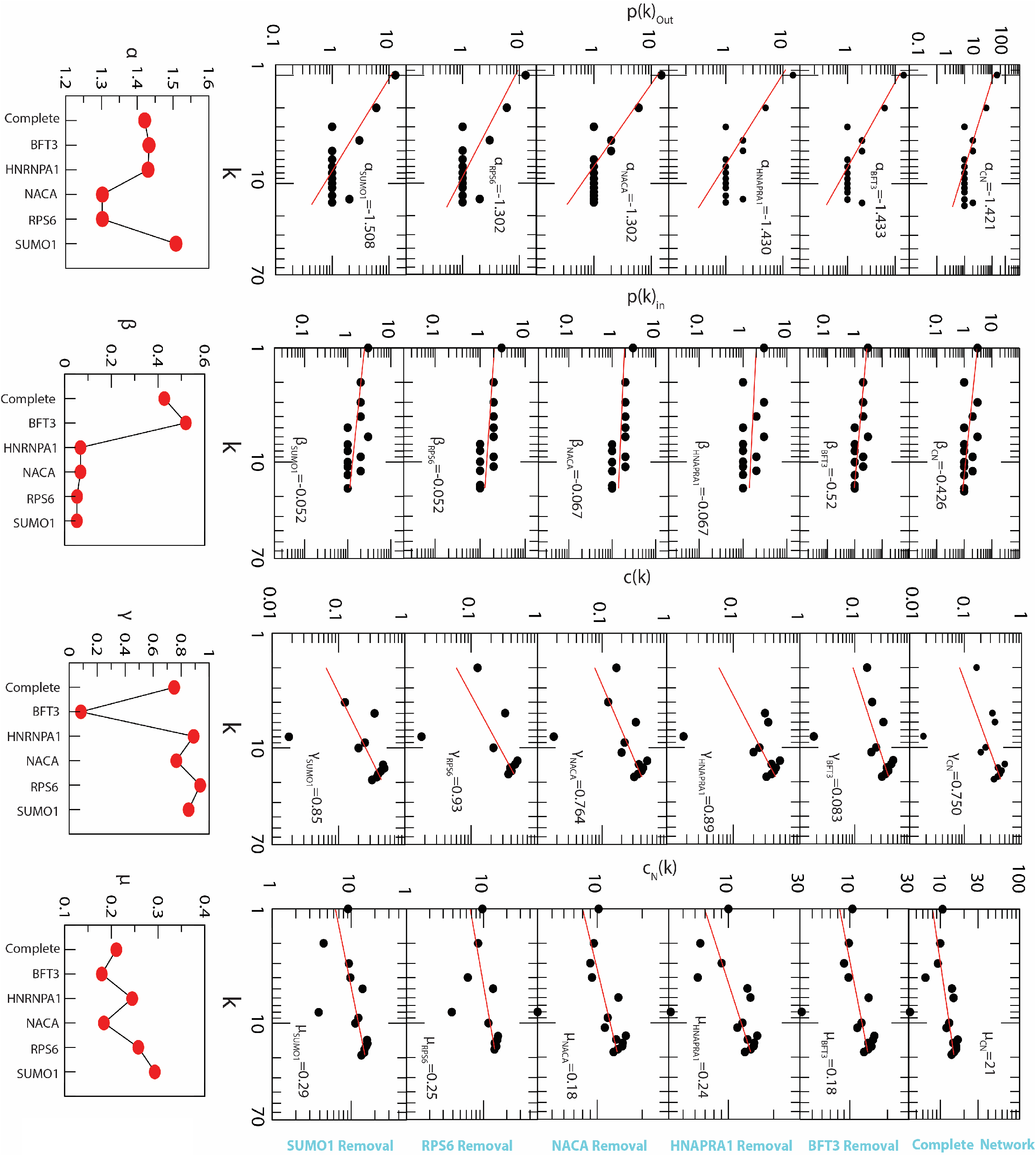
Topological characteristics of the HPIN and hub removal. (a) The figure illustrates the network properties, such as in-degree, out-degree, clustering coefficient, neighbourhood connectivity in the HPIN. The In-degree (P(k)In) and out-degree (P(k)out) distribution, Clustering Co-efficient C(k), Neighbourhood connectivity CN(k) is fitted to the power law distribution with exponent values (α, β, γ & μ). (b) Power law distribution with exponent values α, β, γ & μ of HPIN (complete), hub removed proteins network (RPS6, NACA, HNRNPA1 and BTF3).

## Notes

https://covid19.who.int/

https://gtexportal.org/

https://pubs.broadinstitute.org/mammals/haploreg/haploreg.php

https://www.proteinatlas.org

https://gnomad.broadinstitute.org

https://www.ncbi.nlm.nih.gov/snp.

https://genome.ucsc.edu/

https://www.ncbi.nlm.nih.gov/variation/tools/1000genomes

https://www.datawrapper.de/

https://ldlink.nci.nih.gov/

### Summary of Updates

1. Made corrections to Ms. 2. divided content into main and supplementary data. 3. Included an additional figure as Figure 1' explaining more in a) and b) The findings for the first time, based on potential structural mechanisms, highlighting with ACE2 homodimerisation only one SARS-CoV2 spike protein can bind and further provides explanation how ACE2 can give protection from severe lung injury in COVID19.

