## Supplementary material for "ACE2 Homo-dimerization, Human Genomic variants and Interaction of Host Proteins Explain High Population Specific Differences in Outcomes of COVID19": ACE2 Homodimerization Affects Binding of SARS-CoV-2 Spike Protein

linked to : https://science.sciencemag.org/content/367/6485/1444/tab-e-letters

### ACE2 Homodimerization Affects Binding of SARS-CoV-2 Spike Protein

Yan *et.al.* 2020 proposed the binding of two SARS-CoV-2 Spike Glycoproteins (SGs) with Angiotensin-converting enzyme 2 (ACE2) homodimer without affecting each other or any structural hindrance (*1*). We appreciate the extensive work done and important findings. However, we believe binding of two SGs to ACE2 homodimer is only possible in modelled version as given, but not naturally. At the same time, the alternate explanation we provide may help find an answer to the question that despite being a receptor for SARS-CoV-2, how can ACE2 still perform its physiological role of protection from severe Lung injury, in some COVID-19 patients (*2*). Alignment of receptor binding domain (RBD) (6M17.pdb), with the prefusion SARS-CoV-2 SG (6VSB.pdb), in the optimal ACE2 binding conformation (**Figure 1a)** with "up CTD1" and "open S1 subunit" (*1, 3*)has shown no steric clash with the ACE2 or another SG, as reported (*1*). We extended the evaluation and noted that both SGs protrude radially outward from ACE2 protease domains (PDs). RBDs are 6 nanometer (nm) apart at PDs and anchorage sites of SGs on virus membrane are approximately at a distance of 27 nm (**Figure 1b)**. However, considering the reported inter spike distance of 13-15 nm (*4*) on the viral shell, given almost similar sizes of SARS-CoV (*4, 5*) and SARS-CoV-2 (*6*), SGs are anticipated to be either parallel or protruding radially outwards from its anchorage site (**Figure 1b)**, contrasting to the findings in modelled structure (*1*). Thus, we propose that structural hinderances caused by ACE2 homodimerization, induced by factors like overexpression (*7*), may result in binding of only one SG to dimerized ACE2. The other unbound ACE2 partner might remain free, for its physiological role and continue to protect from lung injury.

**Figure 1. Complex of homodimer of ACE2 with complete SARS-CoV-2 Spike Glycoprotein (SG)**

a) Modelled complex of homodimer of ACE2 colored as brick red (6M17.pdb), with complete SG in open state coloured as sky blue (6VSB.pdb), by aligning the complete SG with RBD colored as purple blue from 6M17.pdb. b) As modelled, the heads of each of the SGs can bind the dimer ACE2, protruding radially outward with RBD 6nm apart but directed inwards, at anchorage sites on virus membrane, with distance of approximately 27 nm whereas, naturally it is either parallel or protruding radially outwards from its anchorage site with inter spike distance of 13-15 nm (*4*).

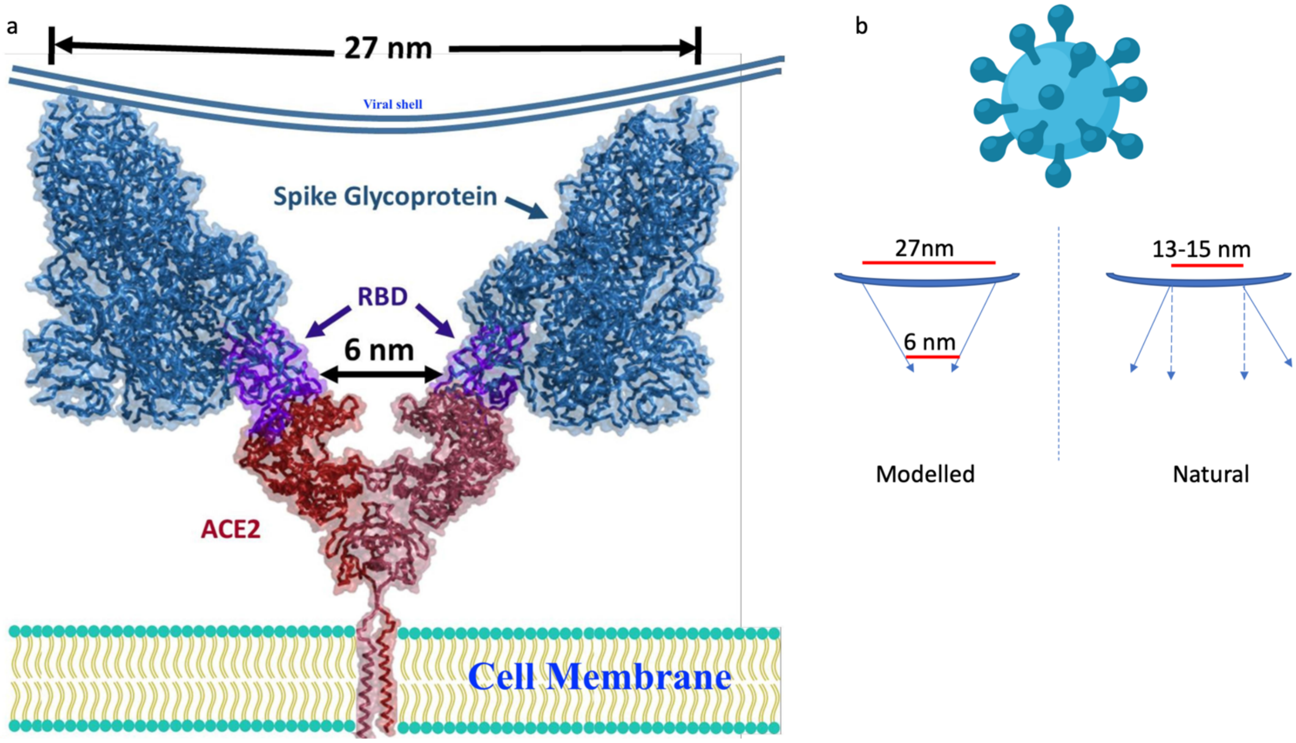
